# Alternative splicing of circadian clock genes correlates with temperature in field-grown sugarcane

**DOI:** 10.1101/718890

**Authors:** Luíza L. B. Dantas, Cristiane P. G. Calixto, Maira M. Dourado, Monalisa S. Carneiro, John W. S. Brown, Carlos T. Hotta

## Abstract

Alternative Splicing (AS) is a mechanism that generates different mature transcripts from precursor mRNAs (pre-mRNAs) of the same gene. In plants, a wide range of physiological and metabolic events are related to AS, as well as fast responses to changes in temperature. AS is present in around 60% of intron-containing genes in Arabidopsis, 46% in rice and 38% in maize and it is widespread among the circadian clock genes. Little is known about how AS influences the circadian clock of C4 plants, like commercial sugarcane, a C4 crop with a complex hybrid genome. This work aims to test if the daily dynamics of AS forms of circadian clock genes are regulated by environmental factors, such as temperature, in the field. A systematic search for AS in five sugarcane clock genes, *ScLHY*, *ScPRR37*, *ScPRR73, ScPRR95* and *ScTOC1* using different organs of sugarcane sampled during winter, with 4 months old plants, and during summer, with 9 months old plants, revealed temperature- and organ-dependent expression of at least one alternatively spliced isoform in all genes. Expression of AS isoforms varied according to the season. Our results suggest that AS events in circadian clock genes are correlated with temperature.

## 1. Introduction

During gene expression in a eukaryotic cell, pre-mRNAs undergo splicing to remove introns and join exons in a mature transcript, generating an open reading frame (ORF) for protein synthesis. Splicing is largely co-transcriptional in yeast, *Drosophila melanogaster*, mammals and *Arabidopsis thaliana* (L.) Heynth. (Beyer and Osheim, 1988; Oesterreich et al., 2016; Saldi et al., 2016; Godoy Herz et al., 2019; Jabre et al., 2019). Alternative splicing (AS) is a mechanism that generates different RNAm transcripts from a single gene. As a result of AS, the mature mRNAs represent another level of gene expression regulation at the post-transcriptional level by, for example, insertion of premature termination codons (PTC), which can target some AS isoforms to degradation by the nonsense-mediated mRNA decay (NMD) pathway (Filichkin and Mockler, 2012; Kalyna et al., 2012; Marquez et al., 2012). Alternatively, those transcripts carrying PTCs could produce truncated polypeptides missing functional domains and motifs that can compete with the corresponding functional protein (Mastrangelo et al., 2012; Reddy et al., 2013; Seo et al., 2011). In addition, AS can increase protein by producing mRNAs from the same gene that encode protein variants with different diversity in function, localization, and stability (Chaudhary et al., 2019; Syed et al., 2012). AS is a ubiquitous process observed from *Drosophila* to humans and plants (Filichkin et al., 2010; Graveley, 2005; Kornblihtt et al., 2013; Marquez et al., 2012). In plants, a wide range of physiological and metabolic events and responses are related to AS. This mechanism is so widespread that it was reported in more than 60% of intron-containing genes in Arabidopsis, 46% in rice and 38% in maize (Chamala et al., 2015; Filichkin et al., 2015c; Marquez et al., 2012; Min et al., 2015; Staiger and Brown, 2013a; Thatcher et al., 2014; Zhang et al., 2010). There is evidence of organ and tissue-specific alternative transcript forms and even alternative transcript isoforms in different subcellular locations (Kriechbaumer et al., 2012; Nagashima et al., 2011; Remy et al., 2013; Vaneechoutte et al., 2017). AS impacts development, from the early gametic cell specification to the seed maturation (Fouquet et al., 2011; Liu et al., 2009; Moll et al., 2008; Sugliani et al., 2010; Szakonyi and Duque, 2018; Zhang et al., 2016) and even flowering time and floral development (Rosloski et al., 2013; Severing et al., 2012; Zhang et al., 2011). Both biotic and abiotic stress responses are also closely related to AS (Shang et al., 2017a; Staiger and Brown, 2013a; Wang et al., 2018). Plants under stress conditions change their AS patterns dramatically (Calixto et al., 2018, 2019; Ding et al., 2014; Filichkin et al., 2015a; Palusa et al., 2007; Staiger and Brown, 2013a). Also, many circadian clock genes generate alternative transcript forms with PTCs under different environmental conditions (Calixto et al., 2016; Filichkin et al., 2010, 2015b; James et al., 2012c, 2012a; Jones et al., 2012). The presence of alternative transcripts in the circadian clock genes is highly conserved among different plant species, such as Arabidopsis, *Populus alba* L., *Brachypodium distachyon* (L.) P. Beauv., and rice (*Oryza sativa* L.) – all C3 plants (Filichkin and Mockler, 2012). Little is known about how AS influences the circadian clock of C4 plants.

The circadian clock is a 24 h endogenous timekeeper mechanism that anticipates the Earth’s day/night and seasonal cycles (Hsu and Harmer, 2014; McClung, 2019; Millar, 2016). Like AS, the circadian clock is associated with growth, photosynthesis, and biomass in plants, so these two regulatory mechanisms may act together, or even regulate each other (Dodd et al., 2005; Harmer, 2009; Lai et al., 2012; Lu et al., 2005; Staiger and Brown, 2013a; Syed et al., 2012). The circadian clock consists of multiple interlocked transcription-translation feedback loops connected with input pathways that feed the circadian clock function with environmental cues, such as light and temperature, and with output pathways that are responsible for coordinating several major metabolic and physiological processes (Haydon et al., 2013; Hsu and Harmer, 2014; Pokhilko et al., 2012). In Arabidopsis, the main loop consists in three different components: *CIRCADIAN CLOCK ASSOCIATED* 1 (*CCA1*), *LATE ELONGATED HYPOCOTYL* (*LHY*), expressed around dawn and *TIMING OF CHLOROPHYLL A/B BINDING PROTEIN 1* (*TOC1*), expressed around dusk (Alabadí et al., 2001). Closely associated with this loop are the *PSEUDO-RESPONSE REGULATORS 7*, *3* and *9* (*PRR7*, *PRR3*, *PRR9*) (Locke et al., 2005; Nakamichi et al., 2010; Para et al., 2007; Zeilinger et al., 2006). The components of the central loop and the associated PRRs are conserved among other plant species, including crops like rice, maize (*Zea mays* L.), barley (*Hordeum vulgare* L.) and sugarcane (*Saccharum* hybrid) (Calixto et al., 2015; Hotta et al., 2013; Khan et al., 2010; Murakami et al., 2007). The sugarcane circadian clock, although sharing conserved components with other plants, may have a broader influence over sugarcane physiology, with 32% of sugarcane transcripts showing rhythms under circadian conditions (Hotta et al., 2013).

Sugarcane is a C4 grass that stores large amounts of sucrose in its stems, which can reach as much as 700 mM or 50% of the culm dry weight (Moore, 1995). Its genome is exceptionally complex, showing aneuploidy and a massive autopolyploidy that can range from six to fourteen copies of each chromosome (Garcia et al., 2013). The genome size of commercial modern sugarcane is estimated to be around 10 Gb (Chan et al., 2018; de Setta et al., 2014). Because modern sugarcane cultivars are interspecific hybrids progenies from *Saccharum officinarum* L. and *Saccharum spontaneum* L., about 80% of sugarcane chromosomes comes from *S. officinarum*, 10% comes from *S. spontaneum* and 10% are recombinants of these two species (Cuadrado et al., 2004; D’Hont, 2005; D’Hont et al., 1996). Sugarcane is a valuable commodity, responsible for 80% of sugar and 40% of ethanol worldwide (FAOSTAT, 2015). The remaining biomass from sugarcane can also be used for bioenergy production: the bagasse can be either burned to generate electricity or have its cell wall hydrolyzed to yield simple sugars, which can be fermented to produce second-generation biofuel (Amorim et al., 2011).

Although a great deal of data has been generated about the plant circadian clock, sugarcane, and AS, the majority of these studies have been performed under highly controlled experimental conditions. Such conditions are essential for reproducibility and, for the circadian clock, a constant environment is one way to demonstrate the inner mechanism generating self-sustained rhythms, as well as rhythmic responses. However, those conditions are far from the environment that crops face in nature, with fluctuations and complex interactions between abiotic and biotic variables (Annunziata et al., 2017, 2018; Shalit-Kaneh et al., 2018). In order to better understand the relationship between the circadian clock and AS and how this relationship impacts on crops, it is essential to expand experiments to field conditions. Indeed, essential in-field studies using Arabidopsis (Annunziata et al., 2018; Richards et al., 2012) and rice (Izawa et al., 2011a; Nagano et al., 2012; Sato et al., 2011) show that the complex natural cyclic environment has a broader impact on rhythmic gene expression. So far, no studies have approached the AS profile on circadian clock genes under such conditions.

In this study, we examined whether the daily dynamics of AS forms of circadian clock genes are regulated by environmental factors in the field. We used sugarcane organs extracted from field-grown plants when individuals were 4-months-old, during the Brazilian winter, and 9-months-old, during the Brazilian summer. We investigated the AS profile of sugarcane circadian clock genes in this fluctuating natural environment. Data shows that there is at least one alternatively spliced form for each of the five circadian clock genes analyzed. During winter, when temperatures are lower, alternative transcripts are more highly expressed than in summer, with higher temperatures, which suggests that AS might be related to the fluctuating environmental temperature in the field. The different organs also showed different levels of AS and leaf has most of the diversity in AS events. Collectively, our data suggest temperature correlates with AS in the circadian clock of sugarcane plants grown in a natural environment, possibly as a mechanism t of dynamic adjustment of the circadian clock.

## 2. Material and Methods

### 2.1. Field Conditions and Plant Harvesting

The sugarcane field where the experiment was conducted was located at the Federal University of São Carlos, campus Araras, in São Paulo state, Brazil (22°21′25″ S, 47°23′3″ W, at an altitude of 611 m). The soil of the site was classified as a Typic Eutroferric Red Latosol. Sugarcane tillers from the commercial variety SP80-3280 (*Saccharum* hybrid) were planted in soil in April/2012. Field design had 8 plots (**Figure S1A**). Each plot had 4 rows containing 20 tillers each. Only sugarcane plants from both central lines were used in order to avoid border effects. Sugarcane individuals were randomly picked from two plots in order to avoid variability of both the local environment and individual plants. Data on environmental conditions was acquired from a local weather station (**Figure S1B – C**). Leaves +1 (L1), a source organ and the first fully photosynthetically active leaf in sugarcane, were sampled from the selected individual plants during two different seasons, and therefore different developmental stages. In the first harvest, 4-months-old plants were sampled in August/2012, during winter; in the second harvest, 9-months-old plants were sampled in January/2013, during summer. In winter, dawn was at 6:30, and dusk was at 18:00 (11.5 h day/12.5 h night). In summer, dawn was at 5:45, and dusk was at 19:00 (13.25 h day/10.75 h night). To compare the rhythms of samples harvested in different seasons, the time of harvesting were normalized to a photoperiod of 12 h day/ 12 h night using the following equations: for times during the day: ZT = 12*T*P_d_^−1^, where ZT is the normalized time, T is the time from dawn (in hours), and Pd is the length of the day (in hours); for times during the night: ZT = 12 + 12*(T – P_d_)*P_n_^−1^, where ZT is the normalized time, T is the time from dawn (in hours), Pd is the length of the day (in hours), and Pn is the length of the night (in hours). Because the 9-month-old plants had their culms fully developed, internodes 1 and 2 (I1) and internode 5 (I5) were also sampled. Both internodes are sink tissues with different profiles: internodes 1 and 2 mostly undergo intense cell division and elongation, whereas internode 5 undergoes sucrose storage. For every time point, 9 individuals were randomly selected in the assigned plots and harvested from the culm up. After that, those 9 individuals were separated into three pools of three individuals, each pool formed a biological replicate and then their leaves +1 were extracted. For all harvests, plants were sampled every 2 h for 26 h, starting 2 h before dawn. In total, the time course consisted of 14 time points in each harvest/season. After every time point sampling, a process that took less than 30 mins on average, tissue was immediately frozen in liquid nitrogen.

### 2.2. RNA Extraction

Sugarcane leaves previously frozen in liquid nitrogen were pulverized using dry ice and a grinder. Then, 100 mg of this ground tissue was used for total RNA extractions using Trizol (Life Technologies, Carlsbad, CA, USA), followed by treatment with DNase I (Life Technologies, Carlsbad, CA, USA) and cleaned with RNeasy Plant Mini Kit (QIAGEN, Valencia, CA, USA). The quality and quantity of each RNA sample were checked using an Agilent RNA 6000 Nano Kit Bioanalyzer (Agilent Technologies, Palo Alto, CA, USA). All RNA samples were stored at −80°C.

### 2.3. cDNA Synthesis

cDNA was synthesized using SuperScript III First-Strand Synthesis System for RT-PCR (Life Technologies, Carlsbad, CA, USA) starting from 5 μg of total RNA. For all reactions, both Oligo(dT) and Random Hexamers primers were used. All cDNA samples were stored at −20°C.

### 2.4. PCR Reactions

Primers used in PCR reactions were designed using the software PrimerQuest Tool (IDT) (http://www.idtdna.com/primerquest/home/index). Each pair of primers was gene-specific and amplified fragments ranging from 242 bp to 805 bp (**Table S1**). All PCR reactions were carried out using Go Taq DNA Polymerase (Promega, Madison, WI, USA) and following the manufacturer’s protocol. Briefly, each 20-μL PCR reaction contained 2 μL of template, 10 μM of each primer, 4 μL of 5x Green Go Taq Buffer, 0.15 μL of Go Taq DNA Polymerase, 2 mM of dNTPs. PCR conditions were: an initial step at 94°C for 2 mins, followed by 20 – 30 cycles of 94°C for 15 s, 50°C for 15 s, 72°C for 30 s, followed by a final extension of 72°C for 5 min. PCR reactions using primers amplifying control genes *ScGAPDH* and *ScPP2AA2* were performed for all cDNA samples. Reactions containing negative control using RNA as template and positive control using genomic DNA as template were carried out. All PCR-amplified fragments were analyzed by taking 10 μL of reaction and run on an electrophoresis gel of 1.5% agarose (Life Technologies, Carlsbad, CA, USA) and 1x TBE (50 mM Tris-HCl pH 8, 50 mM Boric Acid, 1mM EDTA).

**Table 1.**
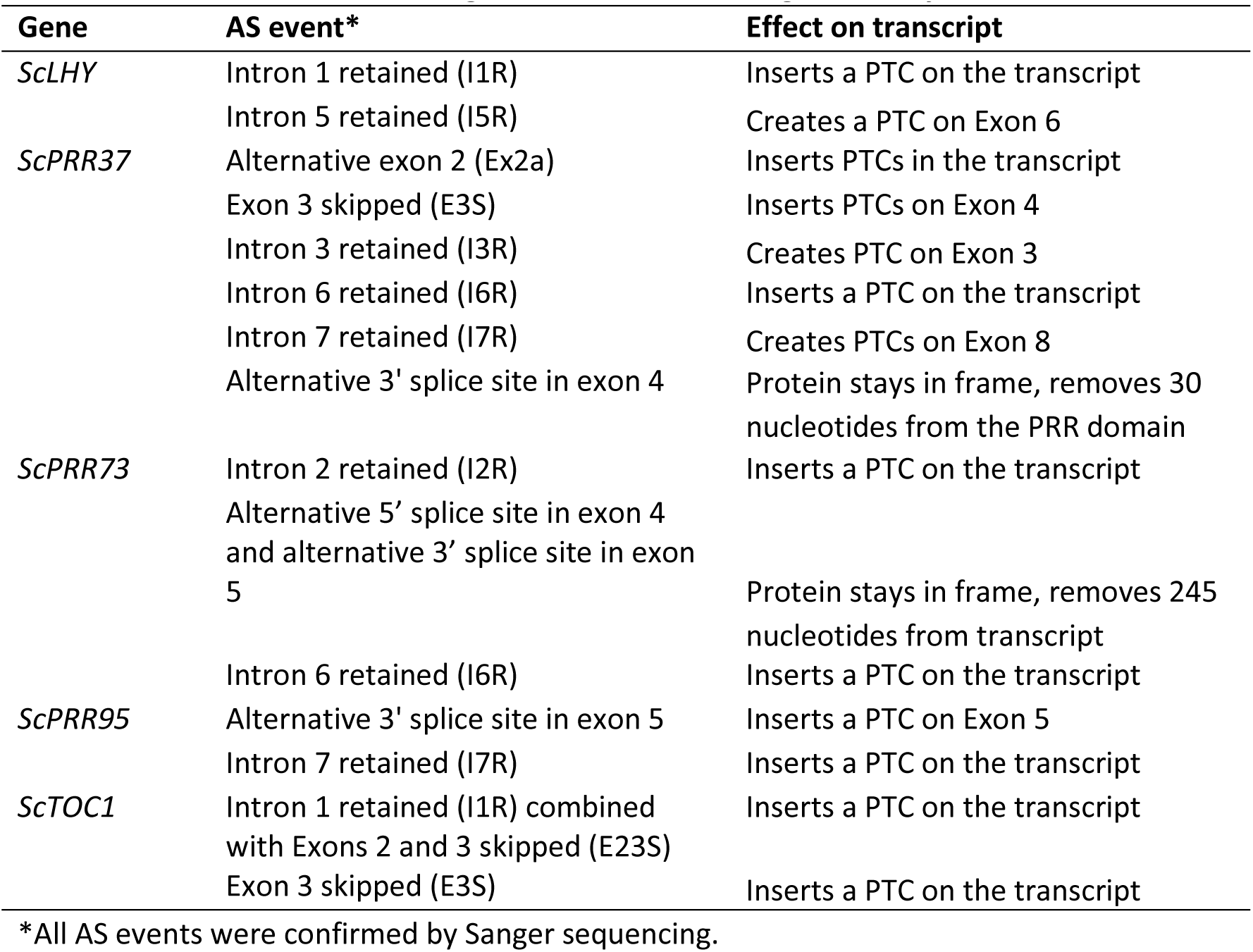
AS events found in the sugarcane circadian clock genes analyzed

### 2.5 High-Resolution RT-PCR

High-Resolution RT-PCR (HR RT-PCR) reactions were performed based on Simpson et al. (2008) and Simpson et al. (2019). For all reactions, the forward primer was labeled with 6-carboxyfluorescein (FAM). Reactions consisted of a final volume of 20 μL which had 2 μL of cDNA, 10 μM of each primer, 2 μL of 10x PCR Reaction Buffer with MgCl2 (Roche Life Science, Indianapolis, IN, USA), 0.15 μL Taq DNA Polymerase (Roche Life Science, Indianapolis, IN, USA), and 2 mM of dNTPs. The PCR detailed program was: an initial step at 94°C for 2 min, followed by 22 – 26 cycles of 94 °C for 15 s, 50°C for 15 s, 70°C for 30 s, followed by a final extension of 70°C for 5 mins. Once PCR reactions were complete, 1 μL of each reaction was added to a mix containing 9 μL of Hi-Di Formamide (Applied Biosystems, Life Technologies, Carlsbad, CA, USA) and 0.5 μL of GeneScan 500 LIZ Size Standard (Applied Biosystems, Carlsbad, CA, USA). The RT-PCR products were separated on an ABI 3730 Automatic DNA Sequencer (Applied Biosystems, Life Technologies, Carlsbad, CA, USA). The results were analyzed using GeneMapper fragment analysis software (Applied Biosystems, Carlsbad, CA, USA). LOESS (locally estimated scatterplot smoothing) regression was used to detect trends in the data. The maximum value of the LOESS curve between ZT0 and ZT22 was considered the peak of the rhythm. The code to fully reproduce our analysis is available on GitHub (https://github.com/LabHotta/AlternativeSplicing) and archived on Zenodo (http://doi.org/10.5281/zenodo.3509232).

### 2.6 Cloning and Sequencing

In order to identify alternatively spliced forms, as well as differentially expressed alleles, RT-PCR fragments were cloned and sequenced. For this, PCR fragments were purified using the QIAquick PCR Purification Kit (QIAGEN, Valencia, CA, USA). Each purified fragment was cloned into pGEM-T Easy Vector (Promega, Madison, WI, USA) following the manufacturer’s protocol. Briefly, each reaction contained 3 μL of purified PCR product, 5 μL of Rapid Ligation Buffer, T4 DNA Ligase, 1 μL of pGEM-T Easy Vector (50 ng) and 1 μL T4 DNA Ligase (3 Weiss units/μL). Ligation reactions were incubated overnight at 4°C. 2 μL of each ligation reactions were used for heat-shock transformation of 50 μL of JM109 High-Efficiency Competent Cells (Promega, Madison, WI, USA), following manufacturer’s instructions. Transformed cells were plated on LB/ampicillin/IPTG/X-gal media and incubated overnight at 37°C. Random colonies were selected to use in plasmid extraction using QIAprep Miniprep Kit (QIAGEN, Valencia, CA, USA). Positive plasmids were confirmed by PCR reactions, digestions using restriction enzymes *PstI* and *Nco1* (Promega, Madison, WI, USA) and Sanger sequencing. In order to identify alternative splicing events and single-nucleotide polymorphisms, results were compared to sugarcane genomic sequences, and sugarcane transcripts from Sugarcane Assembled Sequences (SAS) from SUCEST (http://sucest-fun.org/) using Clustal Omega (https://www.ebi.ac.uk/Tools/msa/clustalo/). *In silico* translation was carried out using ExPaSy translation tool (http://web.expasy.org/translate/).

## 3. Results

### 3.1. Identification of Alternatively Spliced Forms of Sugarcane Circadian Clock Genes

In Arabidopsis, alternative splicing (AS) events of the circadian clock genes are well known (James et al., 2012b). In order to investigate AS of sugarcane circadian clock genes, it is important to first determine and annotate their genomic sequences/structures. We first described the gene structure of previously identified homologs *ScLHY*, *ScPRR73*, *ScPRR95* and *ScTOC1* (Hotta et al., 2013). These homologs were identified by Hotta et al. (2013) using BLAST searches of Arabidopsis and rice circadian clock coding DNA sequence (CDS), on the sugarcane Expressed Sequence Tags (ESTs) collection database at SUCEST (http://sucest-fun.org/). This study identified only CDS sequences of sugarcane circadian clock genes but not their genomic sequences due to the relatively few genomic sequences publicly available at that time for the sugarcane cultivar of interest, plus a high percentage of incomplete genomic sequences (de Setta et al., 2014; Vicentini et al., 2012). Here we used the SUCEST sequences (Hotta et al., 2013) to search for their genomic sequences in an unpublished sugarcane genomic database (Souza et al., unpublished) We identified genomic contigs for *ScLHY* (previously identified as *ScCCA1*), *ScTOC1*, *ScPRR73* (previously identified as *ScPRR3*) and *ScPRR95* (previously identified as *ScPRR59*) (**Table S2**). A new homolog, *ScPRR37*, was identified here for the first time in sugarcane (**Table S2, Figure 1**). The sugarcane genes were annotated by comparison with genomic sequences of circadian clock homologs from barley and sorghum (Calixto et al., 2015). The exon/intron structures are shown in **Figure 1**; introns contained the canonical GT..AG dinucleotides at the exon-intron boundaries. The putative CDS of each gene analyzed was translated *in silico* in order to confirm intact open reading frames (ORFs).

**Figure 1.**
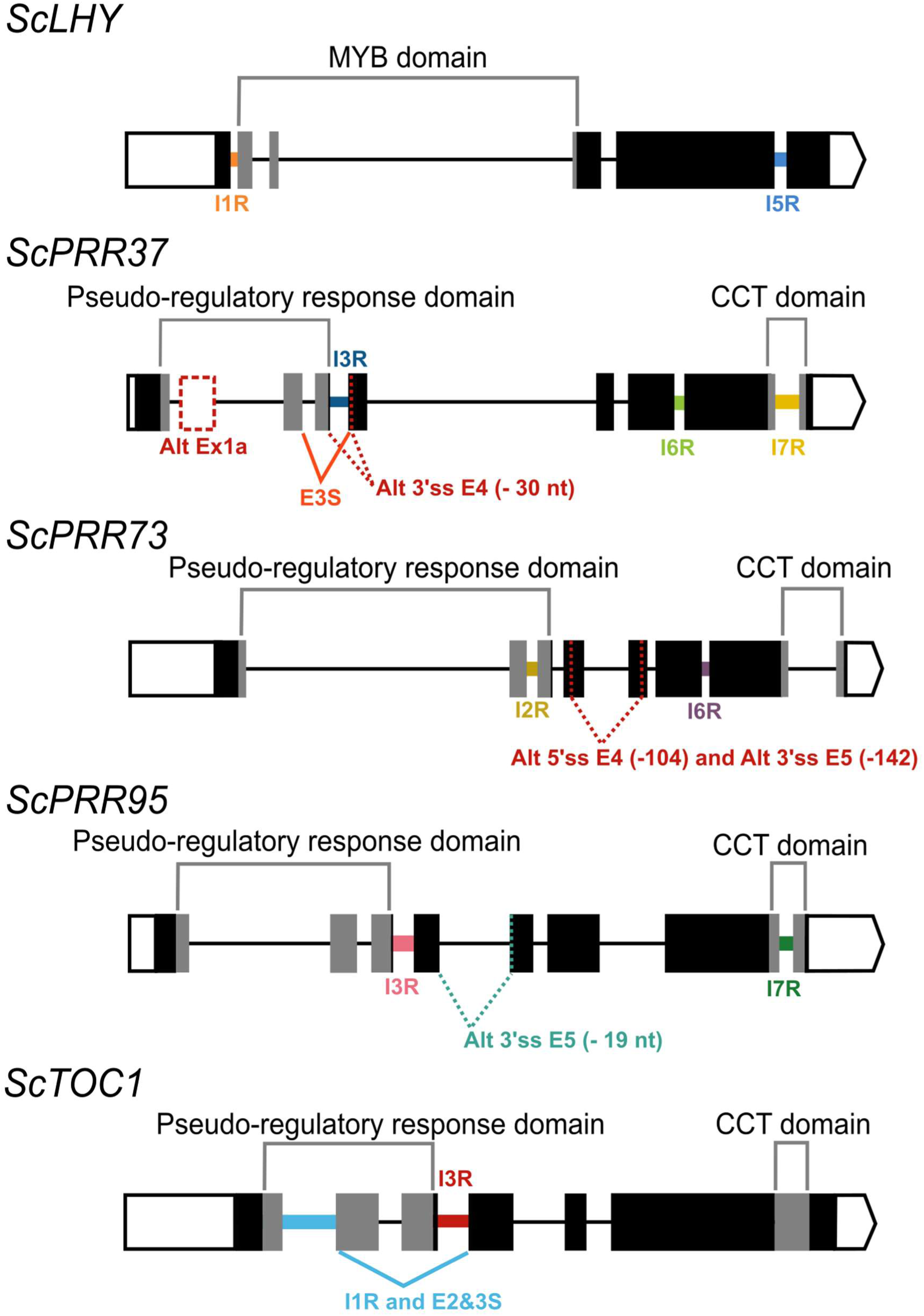
Alternative splicing events identified in the sugarcane circadian clock genes. Alternative splicing (AS) events are shown in the gene structure of *ScLHY, ScPRR37, ScPRR73, ScPRR95*, and *ScTOC1*. White boxes show 5’ and 3’ UTR regions; black boxes show exons; black lines represent introns. The main protein domains are in grey. AS events are shown in colored solid lines, colored dotted lines or in dotted white boxes and the splicing events are illustrated below each event. Different colors were chosen to mark the AS events investigated later, whereas events that were not investigated further are in dark red. IR = intron retention; ES = exon skipping; Alt 5’ ss = alternative 5’ splice sites; Alt 3’ ss = alternative 3’ splice sites; Alt Ex = alternative exon.

To identify AS transcripts, RT-PCR was performed on cDNA synthesized from L1 RNA, collected at three different time points, which corresponded to the highest gene expression for each clock gene, based on the findings from Hotta et al. (2013). RT-PCR used pairs of primers to generate overlapping amplicons to cover the full length of the transcripts for each sugarcane circadian clock genes, *ScLHY*, *ScPRR37*, *ScPRR73*, *ScPRR95*, and *ScTOC1*. RT-PCR products were cloned and sequenced, which confirmed the annotation and identified AS events. Confirmation of different transcript isoforms was carried out by using two different approaches combined: RT-PCR product size and fragment cloning followed by sequencing.

We found evidence of AS events in all five genes analyzed (**Table 1**). Intron retention (IR) was the most frequent AS event identified, resulting in the inclusion of premature termination codons (PTC) in all cases (**Figure 1**). All five genes had at least two IR events, and *ScTOC1* had retention of the first intron (I1R) combined with the skipping of exons 2 and 3 (E23S). *ScLHY* had retention of introns 1 (I1R) and 5 (I5R). *ScPRR37* had retention of introns 3 (I3R), 6 (I6R) and 7 (I7R). *ScPRR73* had retention of introns 2 (I2R) and 6 (I6R). *ScPRR95* had two introns retained: intron 3 (I3R) and intron 7 (I7R) (**Figure 1**). All the other introns in the genes were efficiently spliced. Exon skipping (ES) was detected in *ScPRR37* (E3S) and *ScTOC1*, the latter having two exons skipped at once (E23S), combined with I1R (**Figure 1**). Alternative 3’ splice sites (Alt 3’ ss) were found in both *ScPRR37* and *ScPRR95* in exon 4 (E4) and exon 5 (E5), respectively (**Figure 1**). In *ScPRR73*, we found an AS event between exons 4 and 5 involving an alternative 5’ splice site (Alt 5’ss) in exon 4 (E4) being spliced to an Alt3’ss in exon 5. This AS event removes 104 nt and 142 nt from exons 4 and 5, respectively (total of 246 nt), but remains in frame and potentially produces a protein missing 82 amino acids compared to the wild-type protein.

It is possible that some of the alternative sequences we found are not AS events but different haplotypes since sugarcane is highly polyploid and aneuploid. To exclude this possibility, we obtained the sequences for circadian clock genes from four different sugarcane sequencing projects (Garsmeur et al., 2018; Riaño-Pachón and Mattiello, 2017; Souza et al., unpublished; Zhang et al., 2018). We found 11 sequences for the 5’ portion of the *ScLHY*: 4 from *S. spontaneum* (Zhang et al., 2018), 6 from the commercial *Saccharum* hybrid SP80-3280 (Riaño-Pachón and Mattiello, 2017; Souza et al., unpublished) and 1 from the commercial *Saccharum* hybrid R570 (Garsmeur et al., 2018). All *ScLHY* sequences had the complete intron 1 (**Figure S2**). Similarly, all 16 sequences that had the end portion of *ScLHY* contained the complete intron 5. (**Figure S2**). All other detected AS events had 7 to 15 sequences supporting the conclusion that these alternative sequences are not the result of different sugarcane haplotypes (**Figures S3-S6**).

### 3.2. Expressed AS forms in different seasons in sugarcane leaves

We used HR RT-PCR (Simpson et al., 2019) to examine the daily dynamics of the expressed isoforms of the sugarcane circadian clock genes in two different seasons, winter and summer, using field-grown plants that were 4 and 9 months old, respectively. Briefly, the HR RT-PCR system uses fluorescently labelled primers to amplify across an AS event, followed by fragment analysis in an automatic DNA sequencer that quantifies the relative levels of RT-PCR products and thereby splicing ratios that reflect different splice site choices. The primers used for the HR RT-PCR assays had the same sequence of those used to amplify each gene region on RT-PCR experiments (**Table S1**). Leaf +1 (L1), internode 1 and 2 (I1), and internode5 (I5) samples, harvested every 2 h during 26 h, starting from 2 h before dawn were used. As reference genes to normalize data, we used *ScGAPDH* and *ScPP2AA2* (Iskandar et al., 2004; James et al., 2012a). As the experiments were done in different seasons, we have normalized the time of sampling to fit in a 12 h day/ 12 h night photoperiod such that ZT00 is set to dawn, and ZT12 is set to dusk.

From the five sugarcane circadian clock genes analyzed, HR RT-PCR experiments detected high levels of AS in *ScLHY*, *ScPRR37*, and *ScPRR73* (**Figure 2**), but not in *ScPRR95* and *ScTOC1*, in L1 (**Figure S7**). In general, the AS isoform peaked earlier than the FS form in 8 of the 9 conditions assayed (**Figure 2A-I**), apart from *ScLHY*, where the FS forms peaked at the same normalized time in winter and summer samples.

**Figure 2.**
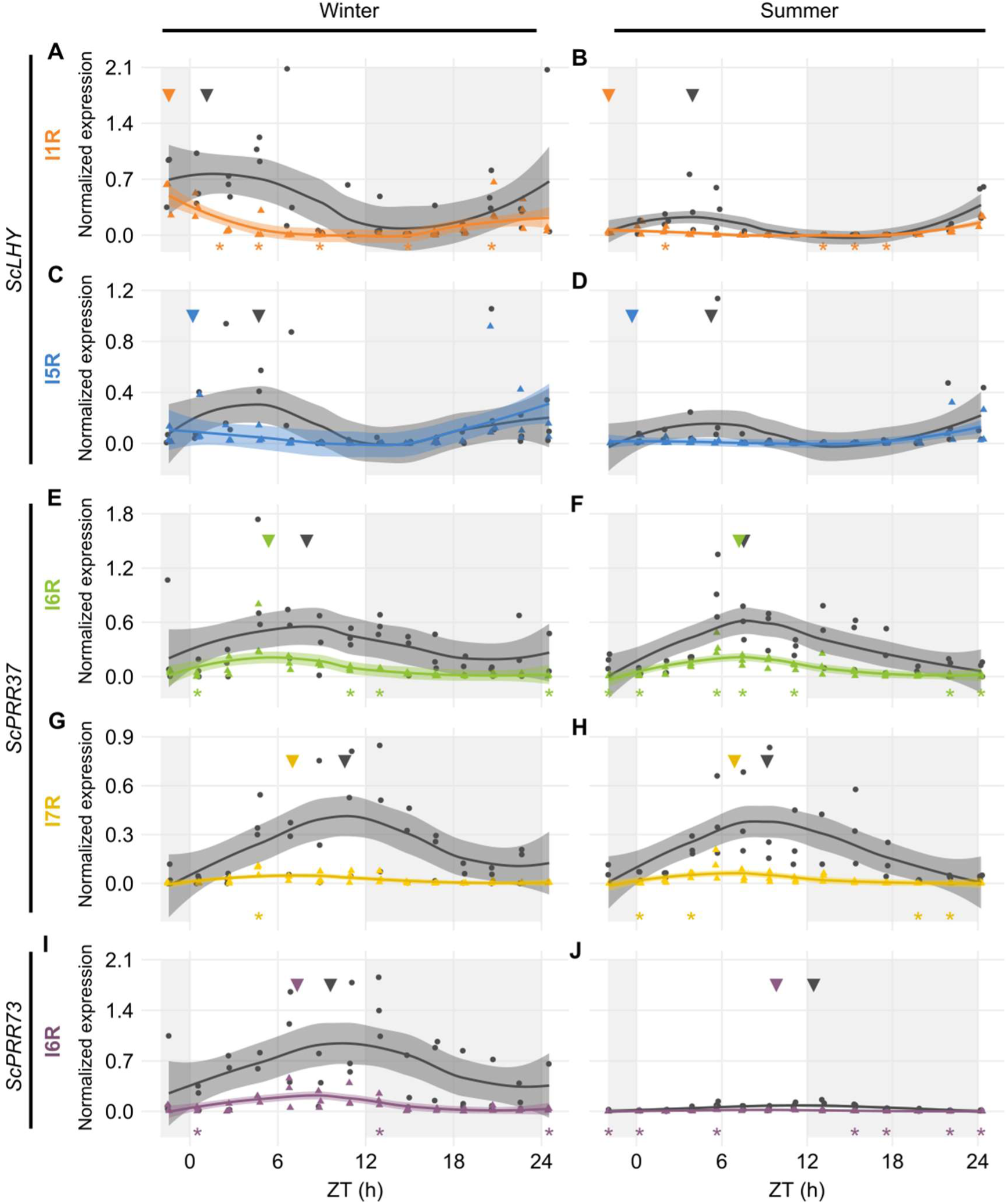
Diel expression profile of fully spliced and alternative transcript isoforms in different seasons. Biological replicates (circles and triangles) and their LOESS curve (continuous lines ± SE) of fully spliced (FS, black) and alternative transcript forms (AS, colored) for the winter samples (4-month-old plants, left) the summer samples (9-month-old plants, right). **(A,B)** *ScLHY* gene expression shows levels of I1R (orange) and **(C,D)** I5R events (blue). **(E,F)** *ScPRR37* gene expression shows levels of I6R (green), and **(G,H)** I6R (yellow); **(I,J)** *ScPRR73* gene expression shows levels of I2R (purple). Inverted triangles show the time of the maximum value of the LOESS curve. The light-gray boxes represent the night period. Statistical significance was analyzed by paired Student’s t-test, * p < 0.05.

*ScLHY* had confirmed events of intron 1 and 5 retention (I1R and I5R) in both harvests (**Figure 2A-D**). The peak of expression of the *ScLHY* alternative isoforms did not match the peak of expression of the fully spliced functional (FS) isoform. *ScLHY* I1R peaked close to dawn in winter and summer samples, while the FS for this region peaked one hour after dawn in winter plants (ZT01) and five hours after dawn (ZT05) in summer plants (**Figure 2A-B**). *ScLHY* I5R peaked at dawn for plants in winter samples, while it was not considered expressed for plants in summer samples (**Figure 2C-D**). The FS form for this region peaked between ZT04-05 for plants in both winter and summer samples (**Figure 2**).

*ScPRR37* I6R had a peak at ZT05 in winter samples, but at ZT07 in summer samples (**Figure 2E and F**). The corresponding FS isoform had a peak at ZT08 in both conditions. In turn, *ScPRR37* I7R had a peak at ZT07 in both winter and summer plants, but the corresponding FS isoform had a peak at ZT090 and ZT11 in winter and summer samples, respectively (**Figure 2G and H**). *ScPRR37* I3R and E3S were not detectable using HR RT-PCR (**Figure S7A-B**). Both the *ScPRR73* I6R and its FS isoform only had high levels in winter samples, with a peak at ZT07 (**Figure 2I-J**). The corresponding FS isoform had a peak at ZT10.

### 3.3. Expressed AS forms in the different source-sink sugarcane organs

We extended the investigation to two other sugarcane organs, internodes 1 and 2 (I1) and internode 5 (I5). These internodes have different physiology: I1 has a high cellular and metabolic activity, whereas I5 is the first internode to actively accumulate sucrose. Only the plants harvested in the summer (9-month-old) had developed internodes that could be harvested. We only measured AS forms in genes *ScLHY* and *ScPRR37*, which were the homologs featuring the highest AS transcript expression in L1 (**Figure 2**). The main difference in expression levels between organs was observed in *ScPRR37* I6R, that had significantly higher levels in leaves during the day compared to the internodes (One-way ANOVA with post-hoc Tukey HSD test, * p < 0.05, **Figure S8**).

In both internodes, *ScLHY* showed detectable levels of both AS events observed in L1. *ScLHY* I1R-containing transcript levels were very low with a peak 2 h before dawn (ZT22), with the FS isoform peaking between ZT03-04 (**Figure 3A-B**). *ScLHY* I5R was also identified in both internodes, at higher levels than *ScLHY* I1R. The AS isoform peaked at ZT22, while the FS form peaked between ZT03-04 (**Figure 3C-D**). During the end of the night, *ScLHY* I1R levels were significantly higher in the leaves compared to the internodes (One-way ANOVA with post-hoc Tukey HSD test, * p < 0.05, Figure S8).

**Figure 3.**
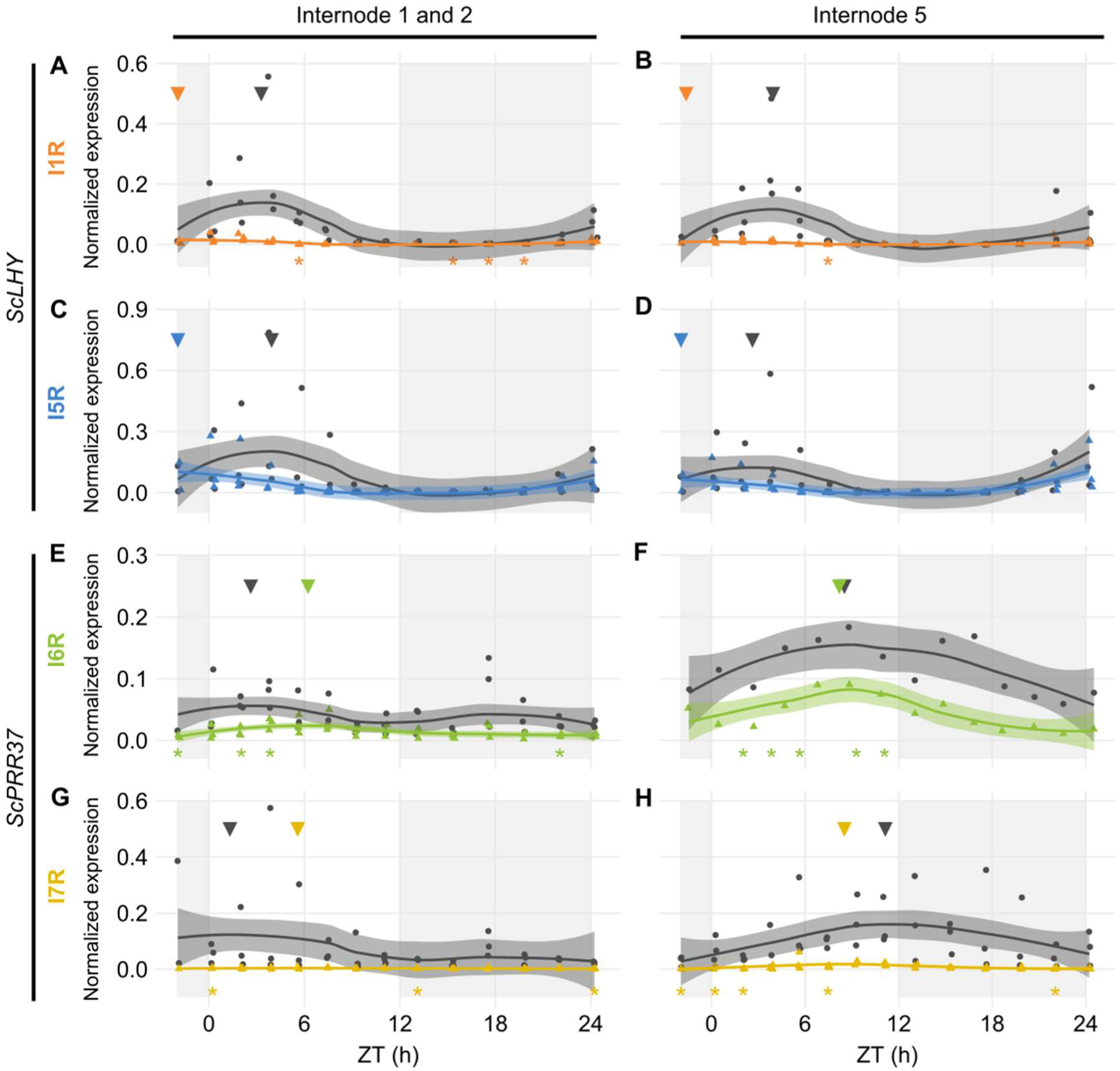
Diel expression profile of fully spliced and alternative spliced transcript isoforms in internodes. Biological replicates (circles and triangles) and their LOESS curve (continuous lines ± SE) of fully spliced (FS, black) and alternative spliced (AS, colored) transcript forms of sugarcane circadian clock genes during the summer harvest in internode 1 and 2 (left) and internode 5 (right). **(A, B)** *ScLHY* gene expression shows levels of I1R (orange) and **(C, D)** I5R (blue); **(E, F)** ScPRR37 gene expression shows levels of I6R (green) and **(G, H)** I7R (yellow). Inverted triangles show the time of the maximum value of the LOESS curve. The light-gray boxes represent the night period. Statistical significance was analyzed by paired Student’s t-test, * p < 0.05.

The *ScPRR37* homolog featured variable levels of FS and AS isoforms when compared to both internodes. In I1, *ScPRR37* I6R and its FS isoform did not have a clear rhythm, with the I6R showing low levels of expression, and the FS isoform showing two peaks. In I5, *ScPRR37* I6R and its FS isoform had higher levels and a peak between ZT08-09 (**Figure 3E-F**). In both internodes, the I7R isoform was expressed at very low levels, but the FS isoform for that transcript region peaked at ZT01 in I1 and ZT11 in I5 (**Figure 3G-H**).

### 3.4. Alternative splicing events are dependent on the time of the day and temperature

After the identification of rhythms in FS and AS transcripts, we tried to identify rhythms in their relative levels by examining the log of the splicing ratio of the AS to the FS transcripts (log(AS/FS)) from the HR RT-PCR data. *ScLHY*, *ScPRR37*, and *ScPRR73* showed evidence of splicing rhythms (**Figure 4**), but only *ScLHY* had more than a 10-fold difference between the maximum and the minimum log(AS/FS) (77-fold) (**Figure 4A**). The AS:FS ratios of all the time-point samples and organs were grouped, as they showed similar rhythmic patterns. In general, all AS events of a gene showed a similar rhythmic pattern. The only exception was E3S in *ScPRR37* (**Figure 4D**), that had a different phase from the other AS events found in *ScPRR37* (**Figure 4B**). The AS events observed in *ScLHY* and E3S in *ScPRR37* had a peak at the end of the night, between ZT20 and ZT24, and a trough between ZT06 and ZT08. In contrast, *ScPRR73* AS events and the remaining *ScPRR37* AS events had a peak between ZT05 and ZT06, and a trough between ZT16 and ZT18.

**Figure 4.**
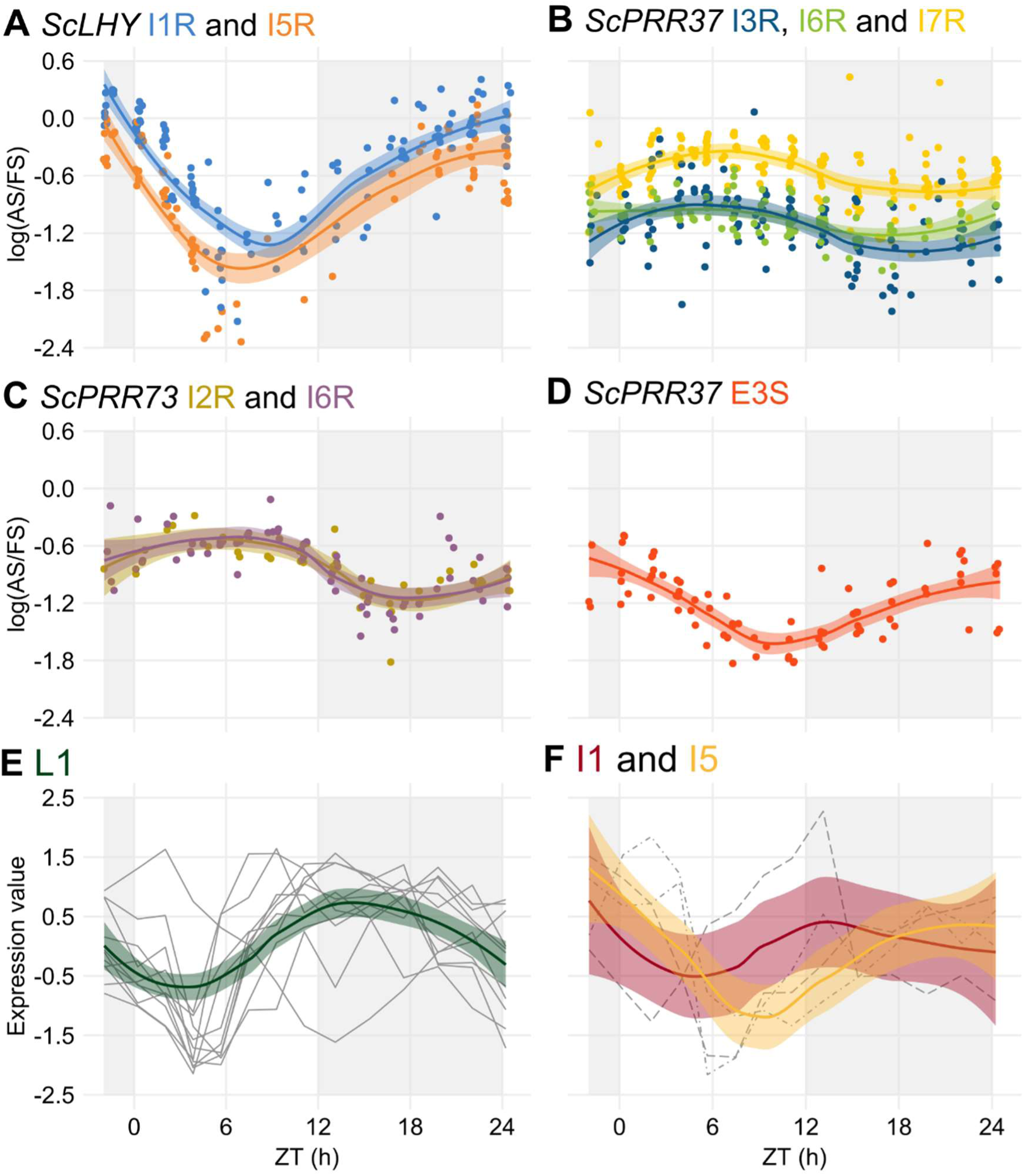
Alternative splicing is rhythmic in sugarcane in field conditions. (**A-D**) The logarithm of the ratio of the expression levels of an AS isoform to its FS isoform, annotated as the log(AS/FS), was plotted against the normalized time of the day (ZT). **(A)** *ScLHY* I1R (orange) and I5R (blue); *ScPRR37* I3R (dark blue), **(B)** I6R (green) and I7R (light yellow); **(C)** *ScPRR73* I2R (gold) and I6R (purple); and **(D)** *ScPRR37* E3S (red). (**E-F**) Normalized expression levels of rhythmic splicing-related transcripts taken from oligo array data (Dantas et al., 2019) in (**E**) leaves +1 (L1, green), and internodes 1 and 2 (I1, red) and (**F**) internode 5 (I5, yellow). Individual expression profiles were drawn in gray. LOESS regression was used to draw the trends in the data in all panels (continuous line ± SE). The light-gray boxes represent the night period.

The rhythmic changes in the log(AS/FS) values could be explained by changes in the expression of putative regulatory genes, such as splicing factors or spliceosomal protein genes. In a previous work, we have identified 6,705 rhythmic transcripts in L1, 3,755 in I1 and 3,242 in I5 in field-grown sugarcane (Dantas et al., 2019). Fourteen spliceosome-related transcripts in the oligo array were expressed in all three organs (**Table S2**): 9 transcripts were rhythmic only in L1, one was rhythmic in L1 and I1, one was rhythmic in L1 and I5, and one was rhythmic in I1 and I5 (**Table S3**). In L1, 9 of the 11 rhythmic transcripts peaked between ZT09 and ZT13, with a trough between ZT03 and ZT5 (**Figure 4E**). In the internodes, most of the transcripts peaked at ZT00 (**Figure 4F**).

To test if the temperature was a factor in the regulation of AS, we correlated temperature information with log(AS/FS) values. Only *ScLHY* AS events showed a significant negative correlation (**Figure 5**). The negative correlation was found in both *ScLHY* AS events, at both harvests/seasons, and all organs. This suggests that the AS regulation of *ScLHY* genes are temperature-dependent.

**Figure 5.**
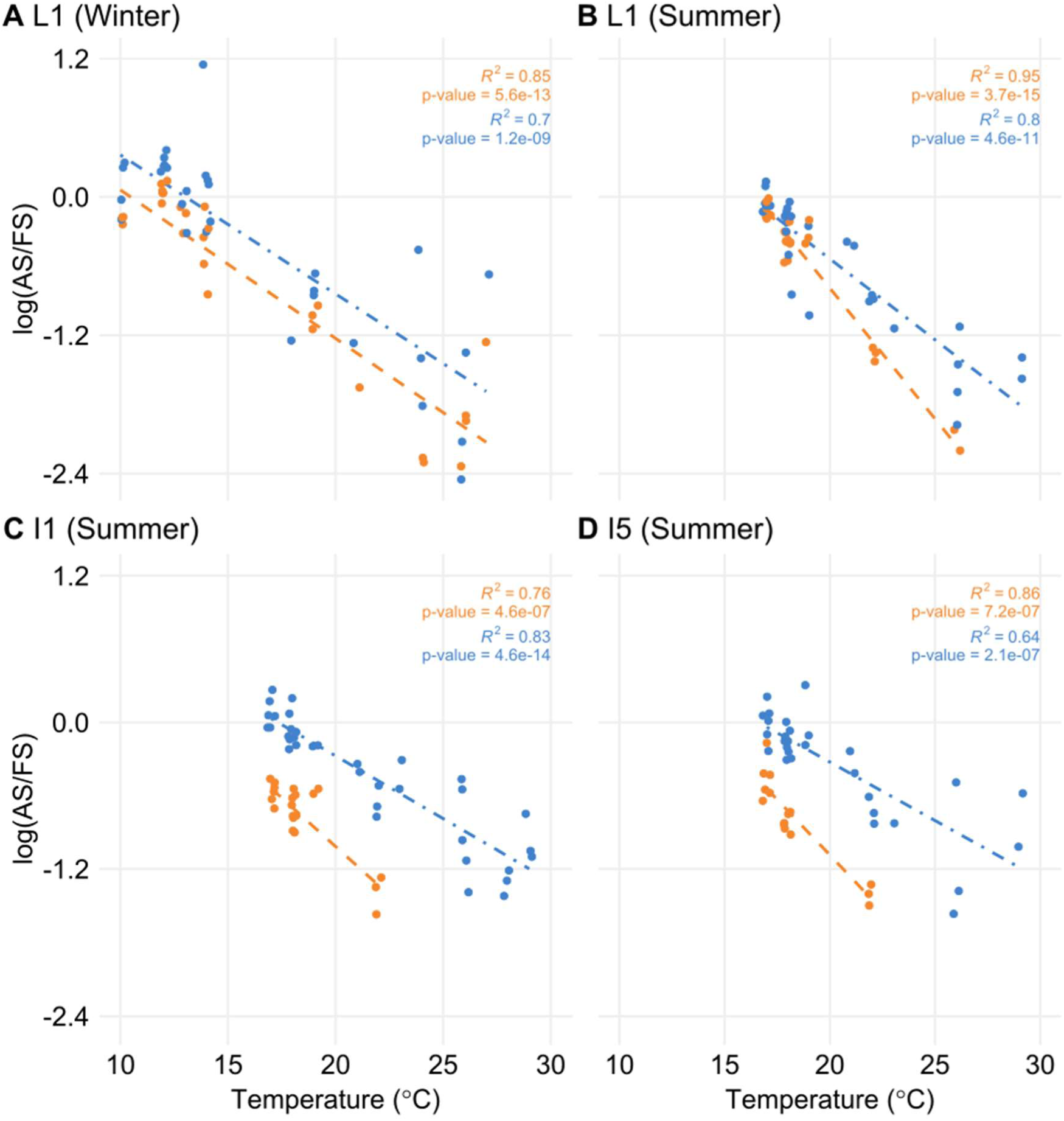
The proportion of alternative spliced and fully spliced forms has a negative correlation with ambient temperature. The logarithm of the ratio of the expression levels of AS and FS isoforms, annotated as the log(AS/FS), was plotted against the ambient temperature for *ScLHY* I1R (orange) and I5R (blue). Regression lines were added for each group of ratios. R^2^ and *P-value* were calculated for each regression. Negative correlations were significant in leaf +1 (L1) in the **(A)** winter (4-month-old plants), and **(B)** summer, **(C)** internode 1 and 2 (I1), and **(D)** internode 5 (I5), both in the summer (9-month-old plants).

## 4. Discussion

In this paper, pioneer information on AS events in the sugarcane circadian clock core genes *ScLHY*, *ScPRR37*, *ScPRR73*, *ScPRR95*, and *ScTOC1* in field-grown sugarcane plants are presented (**Figure 1**). As for Arabidopsis and barley in previous studies, AS is widespread among circadian clock genes (Calixto et al., 2016; Filichkin et al., 2010; Filichkin and Mockler, 2012; James et al., 2012a). The circadian clock homolog *CCA1/LHY* in rice also displays the conserved I1R AS event, suggesting conserved patterns in AS events between the two species (Filichkin et al., 2015b). In barley, AS events were described for the homologs *HvLHY*, *HvPRR37*, *HvPRR73*, and *HvGI*. There are conserved AS isoforms expressed in both Arabidopsis and barley for *HvLHY* and *HvPRR37* (Calixto et al., 2016). The conservation of expression of AS forms across different plant species highlights the role that AS plays in gene expression of circadian clock genes. Our data in sugarcane identified at least one AS event in each of the clock gene homologs analyzed. Given the ploidy of sugarcane, it was necessary to demonstrate that the alternative transcript sequences did not originate from haplotypes but were *bona fide* AS events. By examination of existing sequenced genomic data, we conclude that they are AS events. The most frequent AS event in the sugarcane circadian clock genes was intron retention (IR), which was detected in all five genes. The presence of transcripts containing retained introns might also be an indication of partially spliced transcripts but the other introns in these genes were efficiently removed. The presence of retained introns in transcript isoforms can lead to post-transcriptional regulation of gene expression (Reddy et al., 2013). Since retained introns usually insert premature termination codons (PTCs) in the transcript, they could be substrates for degradation via the nonsense-mediated decay (NMD) pathway. However, in plants, transcripts with detained introns appear to avoid NMD since their abundance is unaffected in NMD mutants (Kalyna et al., 2012; Marquez et al., 2012). Furthermore, such transcripts have been shown to remain in the nucleus and thus avoid the NMD machinery (Göhring et al., 2014). Nuclear intron detention is now recognized as an important post-transcriptional regulatory mechanism (Jacob and Smith, 2017). Intron retention transcripts with PTCs can also potentially give rise to C-terminally truncated and dysfunctional proteins (Mastrangelo et al., 2012; Reddy et al., 2013; Seo et al., 2011). Other AS events in sugarcane circadian clock genes that were in frame and did not insert a PTC in the transcript were the alternative 5’ splice site in ScPRR37 exon 3 (Alt 5’ss E3) and the *ScPRR73* alternative splice site between exons 4 and 5: Alt 5’ss E4 (−104) and Alt 3’ ss E5 (−142) (**Table S2**). In both cases, there is the removal of nucleotides from the coding sequence. *ScPRR37* Alt 5’ss E3 removes 30 nucleotides (10 amino acids) from the PRR domain. The resulting sequence is likely to be translated into a defective protein. In *ScPRR73*, the combination of Alt 5’ss E4 (−104) and Alt 3’ ss E5 (−142) removes 246 nucleotides (82 amino acids) from the coding sequence but still leaves an ORF. However, it is possible that the loss of sequence could affect the normal function of the *ScPRR73* CTT domain. Therefore, the AS events identified in the sugarcane core circadian clock genes either produce transcripts that are likely to be kept in the nucleus and degraded or that are translated into incomplete proteins that are likely to be functionally defective. Thus, AS has an important role in regulating expression and production of core clock proteins. This might have a direct impact on the clock-dependent plant metabolism and physiology.

The levels of alternative transcript isoforms can vary under stress conditions (Calixto et al., 2018; Shang et al., 2017b; Staiger and Brown, 2013b), at specific developmental stages (Szakonyi and Duque, 2018) or in different cell tissues (Shen et al., 2014; Thatcher et al., 2014). In previous work using microarrays, we found that rhythmic expression at the gene level was very organ-specific in sugarcane (Dantas et al., 2019). Our data shows differences in the AS in leaf and internodes at the transcript level. In L1, circadian clock transcripts undergo AS at higher relative levels than in I1 and I5 at the end of the night (**Figure 2 and Figure 3**). In addition, the splicing ratios, log(AS/FS), had rhythms in *ScLHY*, *ScPRR37*, and *ScPRR73*. In *ScPRR37*, the intron retention events peaked in the middle of the day, while the exon skipping event peaked at the end of the night, suggesting that the temporal regulation of these two types of AS events are independent of each other (**Figures 4B and 4D**). Although their expression profiles differ, the consequences of *ScPRR37* I3R and E3S events are likely to be similar. I3R introduces PTCs after exon 3 and E3S removes exon 3 (part of the PRR domain). Arabidopsis *PRR7* also has two mutually exclusive AS events in a similar region of the gene: retention of intron 3 (I3R) and skipping of exon 4 (E4S), both of which give nonproductive mRNAs (James et al., 2012a). The switch from intron retention (mainly during the day) to exon skipping (mainly during the night) in *ScPRR37* may reflect rhythmic changes in specific splicing factors. The splicing ratio rhythms of *ScLHY* and *ScPRR73* showed the same pattern, irrespective of the sampling season or organ (**Figure 4A-C**). This means that these splicing ratios are not organ-specific and are environmentally and circadian clock-regulated in order to have the same distribution during the day and the night regardless of their duration. To try and relate the rhythmic changes in splicing ratios of clock genes to the expression of splicing factor or spliceosomal protein genes, we examined the expression of the splicing-related genes that were rhythmic in L1 in our previous microarray study (**Figure 4E**) (Dantas et al., 2019). The gene-level expression of the majority of these genes peak around dusk (between ZT11 and ZT13), which does not coincide with the peaks in splicing ratios observed in circadian clock genes. The list of splicing-related genes in the microarray analysis was not extensive (**Table S1**) and many splicing factors are alternatively spliced to regulate the level of productive, protein-coding transcripts (Reddy et al., 2013; Staiger and Brown, 2013a). Transcript level RNA-seq will be required to more accurately measure protein-coding transcripts of splicing factors to identify candidate regulators of AS of the core clock genes analyzed here.

Another interesting observation is the noticeable variation in the AS transcripts across the different organs analyzed (**Figure 3**). The differences in transcript expression between source and sink tissues might reflect their metabolical differences. While L1 is a fully photosynthetically active leaf in sugarcane, therefore undergoing photosynthesis, both internodes sink in the assimilated carbon for different purposes: cell division and elongation in I1 and sucrose storage in I5. In *Arabidopsis*, sucrose has been shown to decrease *PRR7* levels, which decrease *ScLHY* levels as a consequence (Frank et al., 2018; Haydon et al., 2013). Thus, differences in *ScPRR37* I6R and *ScLHY* I1R levels (Figure 3 and S8) could be due to the sucrose that is stored in the internodes. In turn, these differences in the levels of circadian clock genes might affect circadian clock outputs. We have found that transcriptional rhythms are very organ-specific in sugarcane (Dantas et al., 2019).

In Arabidopsis, the circadian clock is associated with photosynthesis (Dodd et al., 2009), cell division (Fung-Uceda et al., 2018), and sugar accumulation (Graf et al., 2010; Graf and Smith, 2011; Ko et al., 2016). Taken together, these data suggest that the circadian clock might regulate the different sugarcane organs in distinctive ways. It is already known that metabolites can feedback to regulate the circadian clock, as data show in Arabidopsis that photosynthetic sugars regulate clock functioning (Haydon et al., 2013). Considering that in Arabidopsis different tissues are enriched with different levels of circadian clock transcripts (Endo et al., 2014), a similar phenomenon might occur in sugarcane and explain the different organ transcript expressions and profiles, as well as helping to keep each organ different metabolic and physiologic profiles.

We found that the splicing ratios from *ScLHY* AS transcripts are negatively correlated with temperature in all organs. Previous studies on the presence of AS in circadian clock genes in Arabidopsis and barley revealed that, under controlled conditions, the AS status of circadian clock genes is regulated by temperature, especially *LHY* and its paralog *CCA1* (Calixto et al., 2016, 2018; Filichkin et al., 2015a; James et al., 2012a and b; Kwon et al., 2014; Marshall et al., 2016; Park et al., 2012; Seo et al., 2012). Lower temperatures led to increased abundance of alternative non-productive forms of *LHY*, *PRR7* and *PRR5* in Arabidopsis (James et al., 2012a). This could affect the expression of fully spliced transcripts of the circadian clock genes, promoting a functional modulation in the circadian clock central oscillator, which might be reflected by altered temporal control of the clock outputs. In Arabidopsis, cold temperatures reduced the amplitude of *CCA1*/*LHY*, as well as disrupted the circadian clock function (Bieniawska et al., 2008). Considering the natural environment context, where plants like sugarcane face fluctuations in temperature on a daily and yearly basis, the continuous temperature-regulation of AS of the circadian clock network could have a more profound impact on metabolism and, ultimately, on crop yield.

The data in our work shows that from winter to summer, as the temperature increases (**Figure S1B**), the expression of alternative forms of the circadian clock genes decreases, noticeably for *ScLHY* (**Figure 2A-D**). In Arabidopsis, there is evidence linking the circadian clock with sugar accumulation as starch through *CCA1* and *LHY* (Miller et al., 2012; Ng et al., 2014; Ni et al., 2009) and in field-grown maize, a C4 plant like sugarcane, two *CCA1* homologs are associated to photosynthesis and, therefore, sugar accumulation (Ko et al., 2016). All this evidence allows us to speculate that there might be differences in the sugar accumulation by the field-grown sugarcane from winter to summer, but a metabolomic analysis focused on sucrose and hexoses content would be necessary to bring evidence to support such speculation.

Recent studies featuring experiments conducted in field conditions in Arabidopsis, rice and tomato highlight the differences in gene expression, circadian regulation and plant metabolism compared to experiments conducted inside growth chambers (Annunziata et al., 2017, 2018; Higashi et al., 2016; Izawa et al., 2011b; Shalit-Kaneh et al., 2018). Because AS has an impact on the regulation of gene expression, which impacts circadian regulation and plant metabolism, it is important to start investigating the dynamic adjustment of AS in response to a fluctuating environment. RNA-seq data from time-series experiments revealed that the AS status of Arabidopsis transcriptome is widely responsive to changes in temperature (Calixto et al., 2018). By progressively lowering temperature, rapid changes in the spliced forms of transcripts were detected, which suggests that AS might also act to regulate low-temperature responses and how plants tolerate such stress (Calixto et al., 2018). The Arabidopsis circadian clock also acts in regulating plant abiotic stress tolerance (Grundy et al., 2015). This also suggests that both AS and the circadian clock might act in synergy to help plants to cope with temperature changes in both the short and long term. In the field, this regulation might be even more important, due to the unexpected fluctuations in light, temperature, and humidity in which plants are exposed.

Our data show that AS occurs in sugarcane circadian clock genes and that the different transcript isoforms show a dynamic expression profile in sugarcane grown under field conditions. Furthermore, *ScLHY* AS regulation correlates with temperature in sugarcane circadian clock genes. Thus, the changes in expression of alternative isoforms of *ScLHY* transcripts observed across winter and summer might illustrate the combined effect of both the circadian clock and AS regulation and AS in *ScLHY* might be a key mechanism that allows the continuous dynamic adjustment of the circadian clock by temperature in sugarcane. It is important to start further studies on the impact of the seasonal variation on the AS isoforms of the circadian clock gene expression and, ultimately, sugarcane metabolism and yield.

## 5. Author Contributions

CTH and JWSB designed this research. LLBD and CTH harvested the biological material and carried out BLAST analyses. LLBD processed all samples and carried out cloning, HR RT-PCR and SNP analyzes. LLBD, CPGC, JWSB, and CTH participated in the interpretation of genomic annotation and HR RT-PCR data. MMD contributed with cloning. MSC contributed with the plants and space for the field experiment. LLBD and CTH drafted the manuscript. All authors participated in its correction and have read and approved the final manuscript. CTH and JWSB acquired the funding.

## 7. Acknowledgments

The present study was supported by the São Paulo Research Foundation (FAPESP) [grant nos. 11/00818-8 and 15/06260-0; BIOEN Program], and by the Serrapilheira Institute (grant no. Serra-1708-16001). L.L.B.D. was supported by FAPESP scholarships [grants 11/08897-4 and 15/10220-3]. C.P.G.C and J.W.S.B. were supported by funding from the Biotechnology and Biological Sciences Research Council (BBSRC) [BB/K006568/1 and BB/N022807/1] and the Scottish Government Rural and Environment Science and Analytical Services division (RESAS) [to J.W.S.B.].

## 8. Supplementary Material

The Supplementary Material for this article can be found online at: (future link here)

---

**Supplemental Figure 1.**
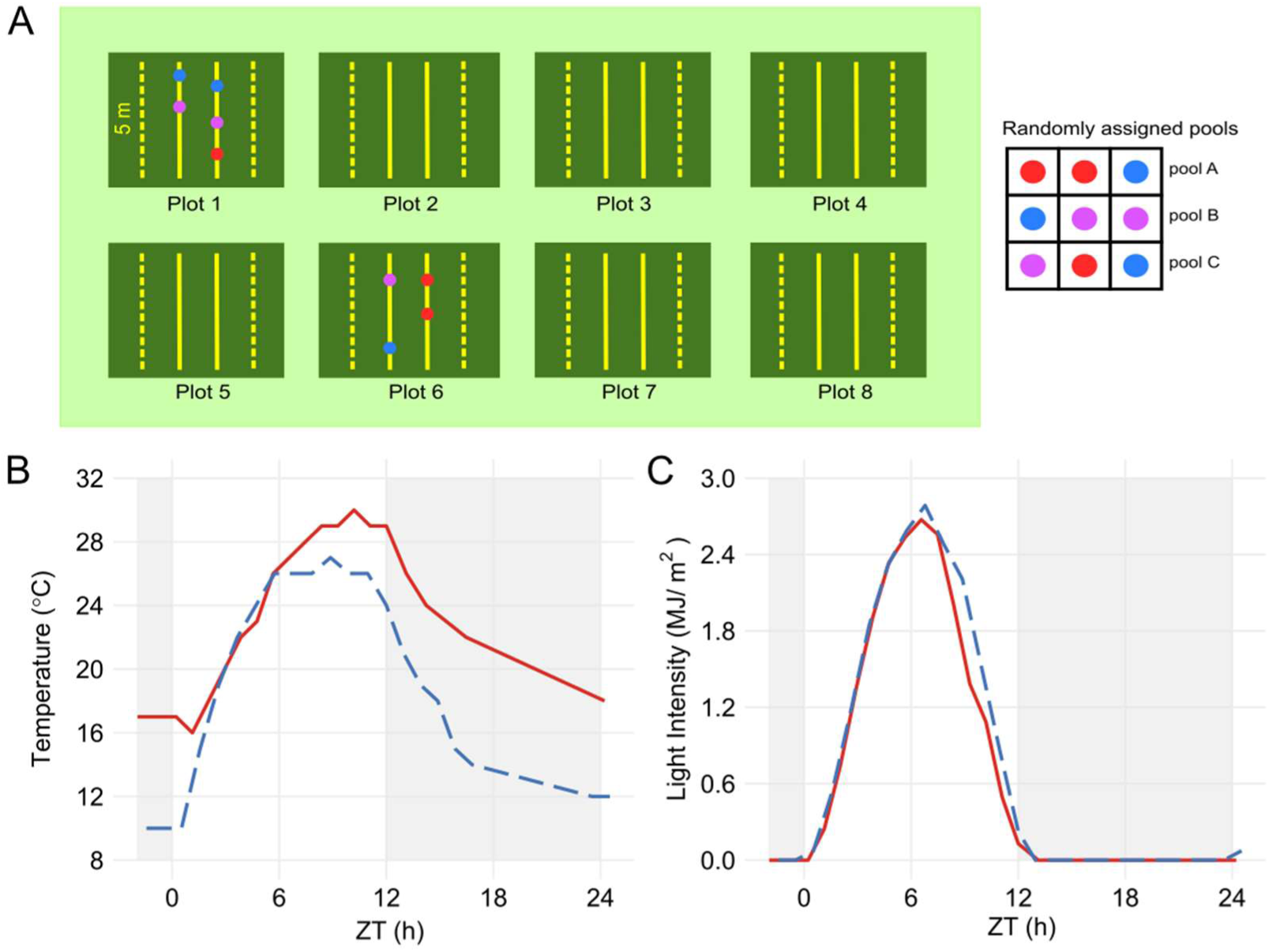
Field design and environmental conditions. **(A)** The experimental field consisted of 8 plots, that were surrounded by two lines of sugarcane. There were 4 lines per plot, each line 5 m long and filled with 20 tillers of commercial sugarcane (variety SP80-3280). The border lines (dashed yellow lines) were not used in our experiment in order to avoid border effects. Plants from the two central lines of two plots were randomly picked and randomly assigned to three pools. Temperature **(B)** and light intensity **(C)** during the summer sampling (red continuous line) and the winter sampling (dashed blue line) were obtained from a nearby weather station. The time points were normalized for a 12h photoperiod. The light-gray boxes represent the night period.

**Supplemental Figure 2.**
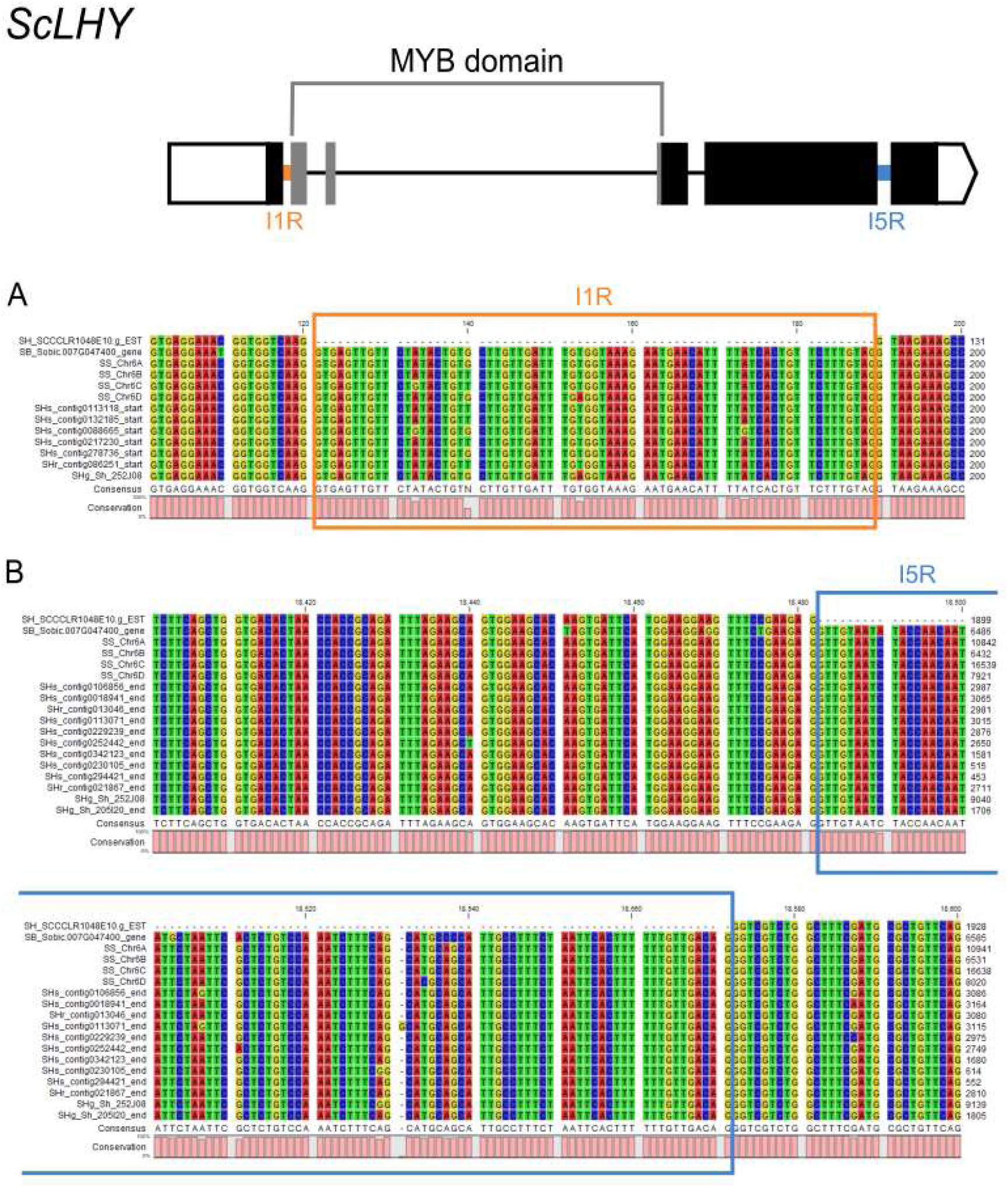
Alignment of two different regions of ScLHY from different genomic sequences. Sequences from different sugarcane genome projects that were identified as *ScLHY* were aligned in order to compare the regions containing the I1R and I5R AS events in different haplotypes. The regions containing (**A**) the intron 1 (orange) and (**B**) intron 5 (blue) were highlighted to show that all the sequenced haplotypes have the same gene structure. Sequences are from *Sorghum bicolor* (SB) genome (McCormick et al., 2018), *S. spontaneum* (SS) (Zhang et al., 2018), *Saccharum* hybrid (SP80-3280): SHs (Souza et al., unpublished), SHr (Matiello et al., 2017), and *Saccharum* hybrid (R570) (SHg) (Garsmeur et al., 2018).

**Supplemental Figure 3.**
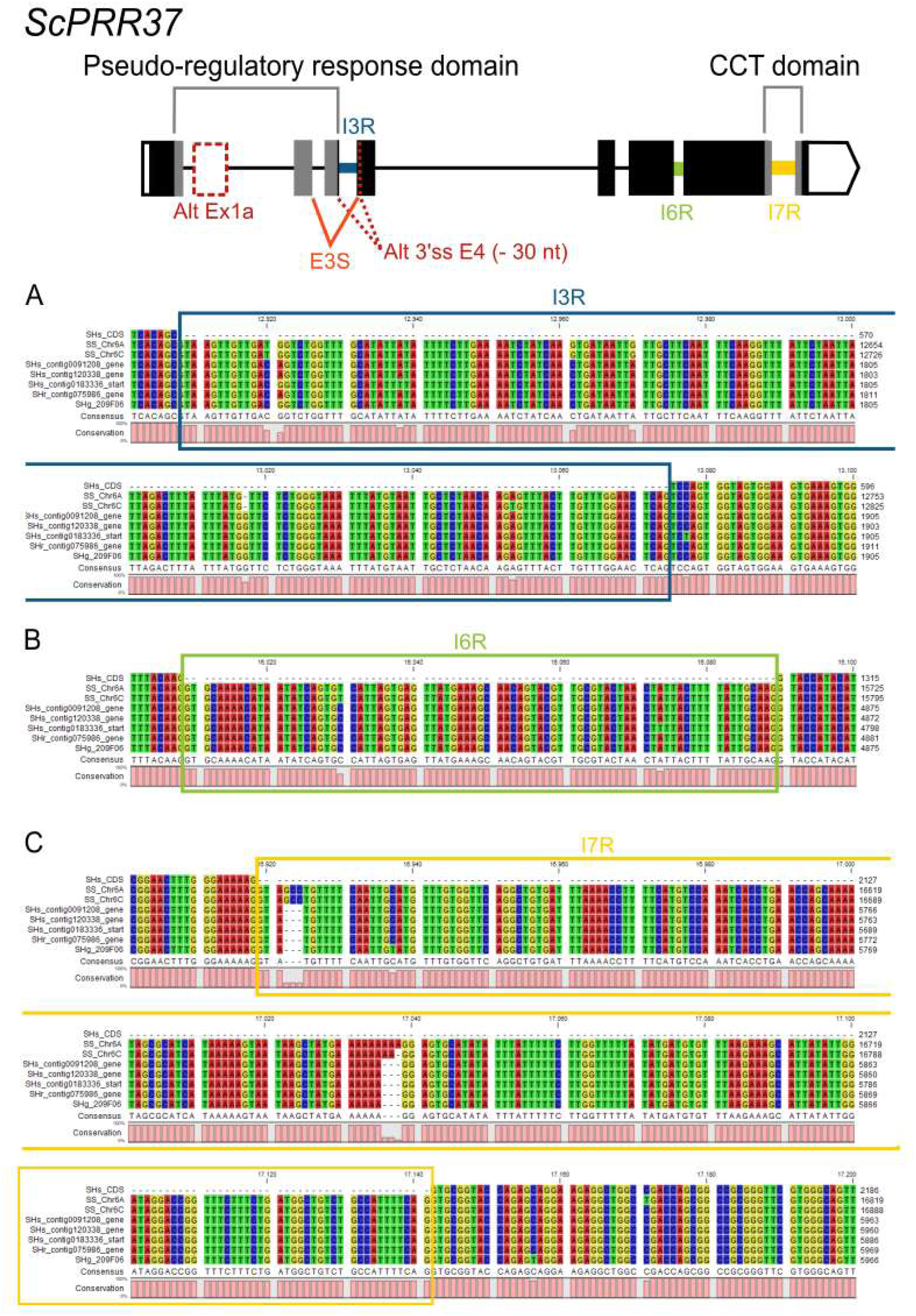
Alignment of three different regions of *ScPRR37* from different genomic sequences. Sequences from different sugarcane genome projects that were identified as *ScPRR37* were aligned in order to compare the regions containing the I3R, I6R, and I7R AS events in different haplotypes. The regions containing (**A**) the intron 3 (dark blue), (**B**) intron 6 (green) and (**C**) intron 7 (yellow) were highlighted to show that all the sequenced haplotypes have the same gene structure. Sequences are from *S. spontaneum* (SS) (Zhang et al., 2018), *Saccharum* hybrid (SP80-3280): SHs (Souza et al., unpublished), SHr (Matiello et al., 2017), and *Saccharum* hybrid (R570) (SHg) (Garsmeur et al., 2018).

**Supplemental Figure 4.**
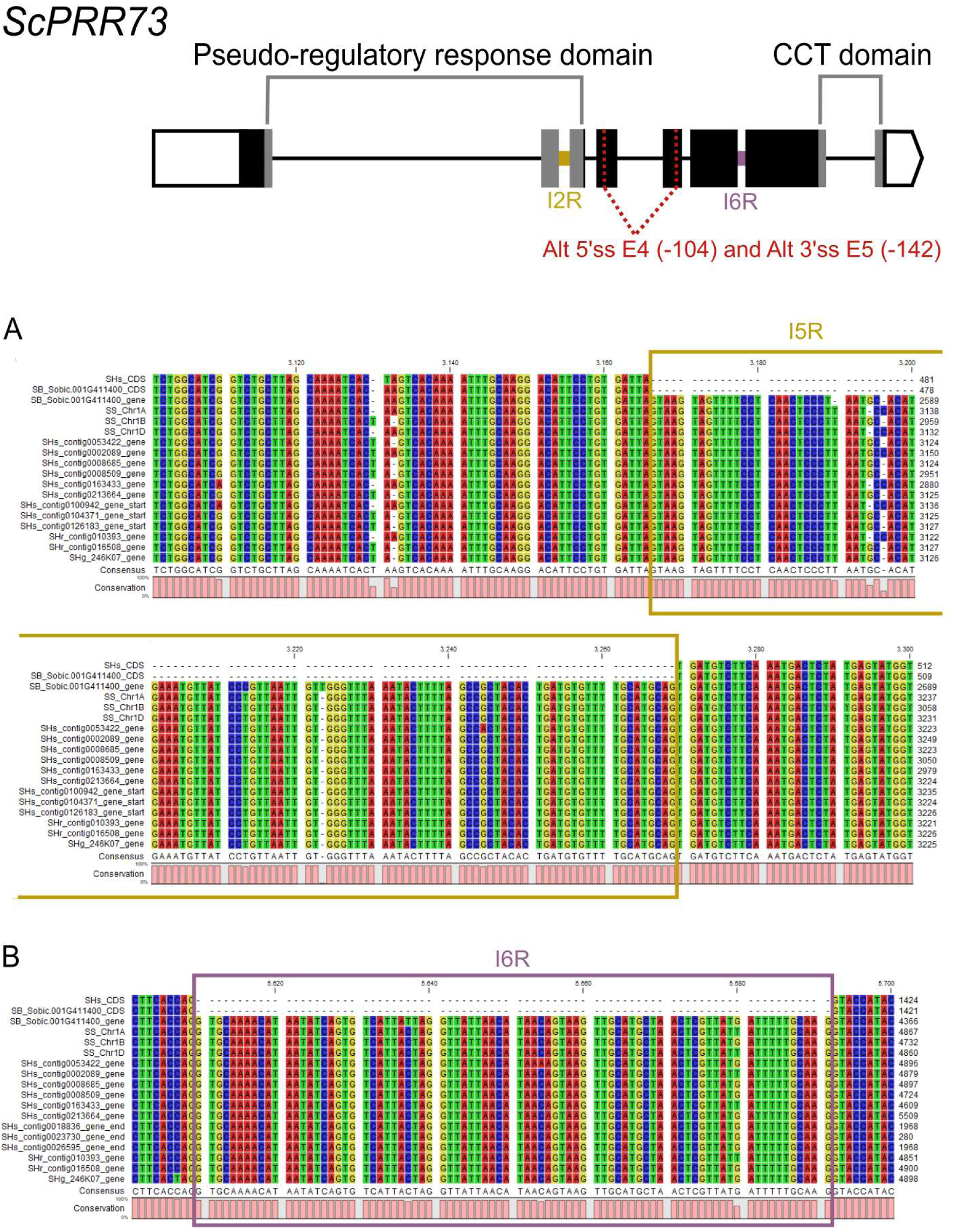
Alignment of two different regions of *ScPRR73* from different genomic sequences. Sequences from different sugarcane genome projects that were identified as *ScPRR73* were aligned in order to compare the regions containing the I2R and I6R AS events in different haplotypes. The regions containing (**A**) the intron 2 (gold) and (**B**) intron 6 (purple) were highlighted to show that all the sequenced haplotypes have the same gene structure. SS sequences are from *S. spontaneum* (Zhang et al., 2018). SHs are from the *Saccharum* hybrid (SP80-3280) sequenced by Souza et al. (unpublished), and SHr were sequenced by Matiello et al. (2017). SHg are from the *Saccharum* hybrid (R570) sequenced by Garsmeur et al. (2018).

**Supplemental Figure 5.**
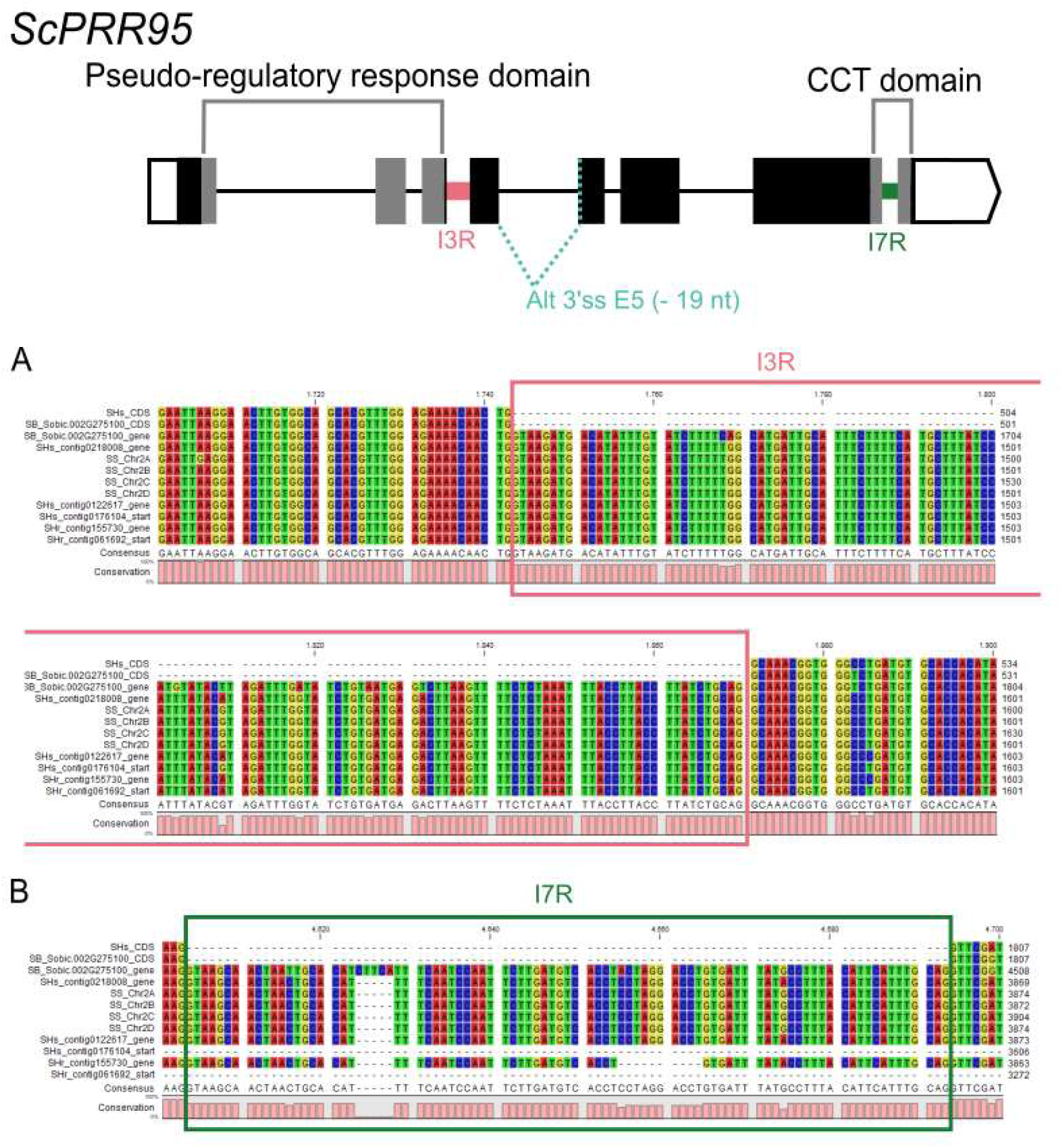
Alignment of two different regions of *ScPRR95* from different genomic sequences. Sequences from different sugarcane genome projects that were identified as *ScPRR95* were aligned in order to compare the regions containing the I3R and I7R AS events in different haplotypes. The regions containing (**A**) the intron 3 (pink) and (**B**) intron 7 (dark green) were highlighted to show that all the sequenced haplotypes have the same gene structure. SS sequences are from *S. spontaneum* (Zhang et al., 2018). SHs are from the *Saccharum* hybrid (SP80-3280) sequenced by Souza et al. (unpublished), and SHr were sequenced by Matiello et al. (2017). SHg are from the *Saccharum* hybrid (R570) sequenced by Garsmeur et al. (2018).

**Supplemental Figure 6.**
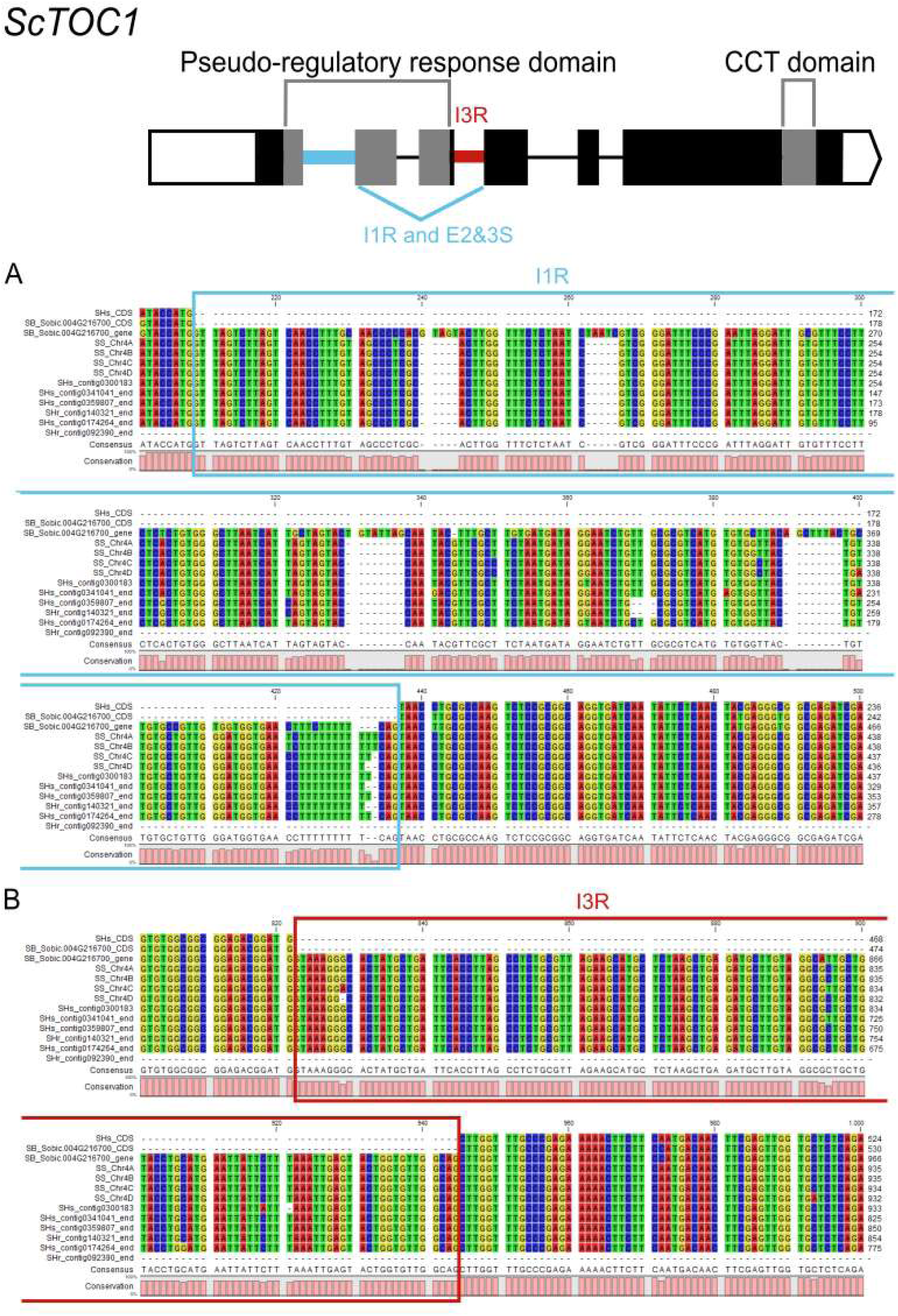
Alignment of two different regions of *ScTOC1* from different genomic sequences. Sequences from different sugarcane genome projects that were identified as *ScTOC1* were aligned in order to compare the regions containing the I1R and I3R AS events in different haplotypes. The regions containing (**A**) the intron 1 (light blue) and (**B**) intron 3 (red) were highlighted to show that all the sequenced haplotypes have the same gene structure. SS sequences are from *S. spontaneum* (Zhang et al., 2018). SHs are from the *Saccharum* hybrid (SP80-3280) sequenced by Souza et al. (unpublished), and SHr were sequenced by Matiello et al. (2017). SHg are from the *Saccharum* hybrid (R570) sequenced by Garsmeur et al. (2018).

**Supplemental Figure 7.**
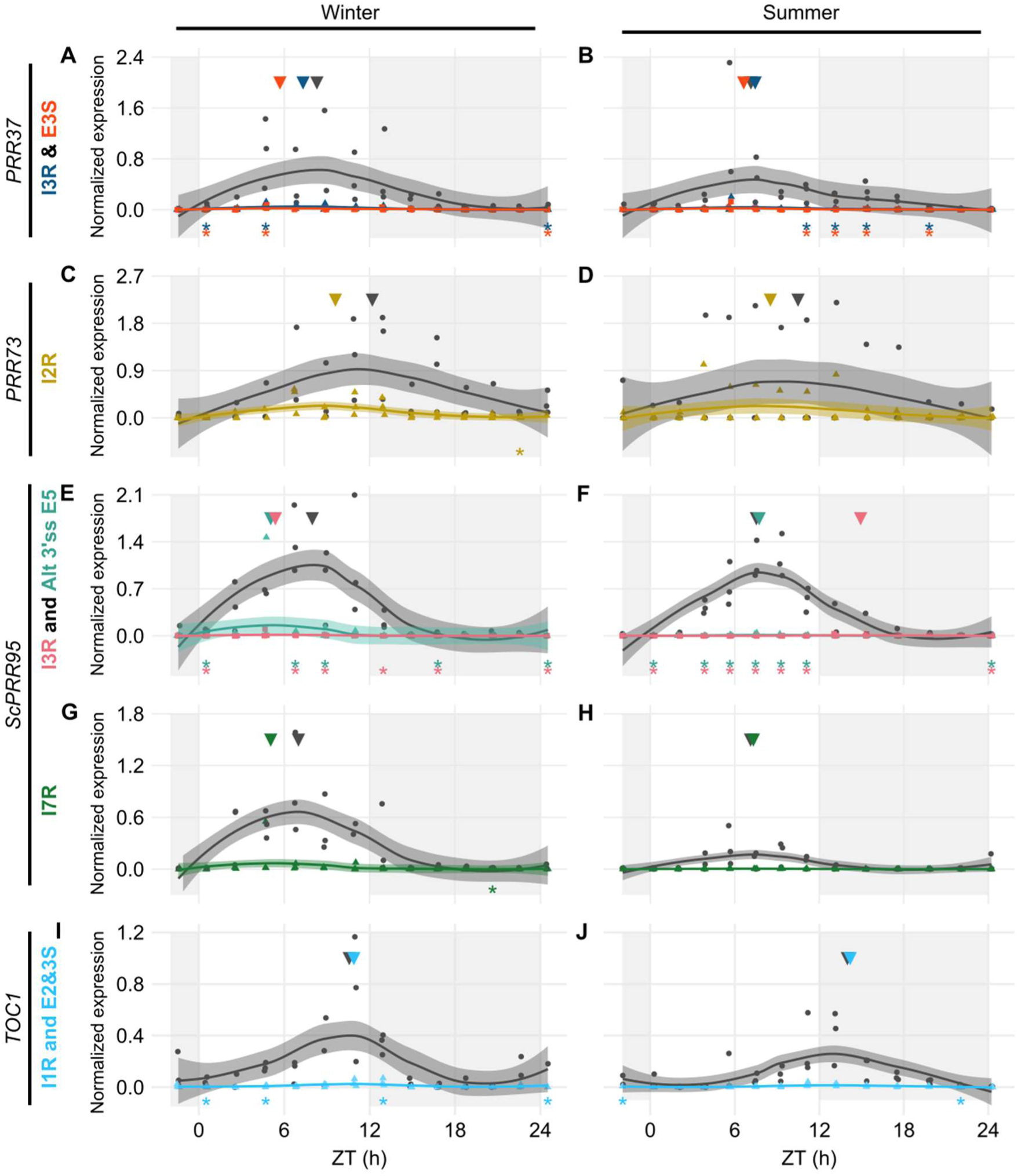
Diel expression profile of fully spliced and alternative transcript isoforms in different seasons. Biological replicates (circles and triangles) and their LOESS curve (continuous lines ± SE) of fully spliced (FS, black) and alternative transcript forms (AS, color) for the winter samples (4-months old plants, left) the summer samples (9-months old plants, right). These AS forms, though detected by HR RT-PCR, were not highly expressed. *ScPRR37* gene expression shows levels of I3R (orange) and E3S (dark blue) (**A-B**); *ScPRR73* gene expression shows levels of I2R (gold) (**C-D**); *ScPRR95* gene expression shows levels of I3R (pink) and Alt 5’ss E5 (teal) (**E-F**), and I7R (dark green) (**G-H**). *ScTOC1* gene expression shows levels of I1R/E23S (cyan) (**I-J**). Inverted triangles show the time of the maximum value of the LOESS curve. The light-gray boxes represent the night period. Statistical significance was analyzed by paired Student’s t-test, * p < 0.05 (C-D, G-J) or by One-way ANOVA with post-hoc Tukey HSD test, * p < 0.05 (A-B, E-F), each color corresponding to a test of an AS form against the FS form.

**Supplemental Figure 8.**
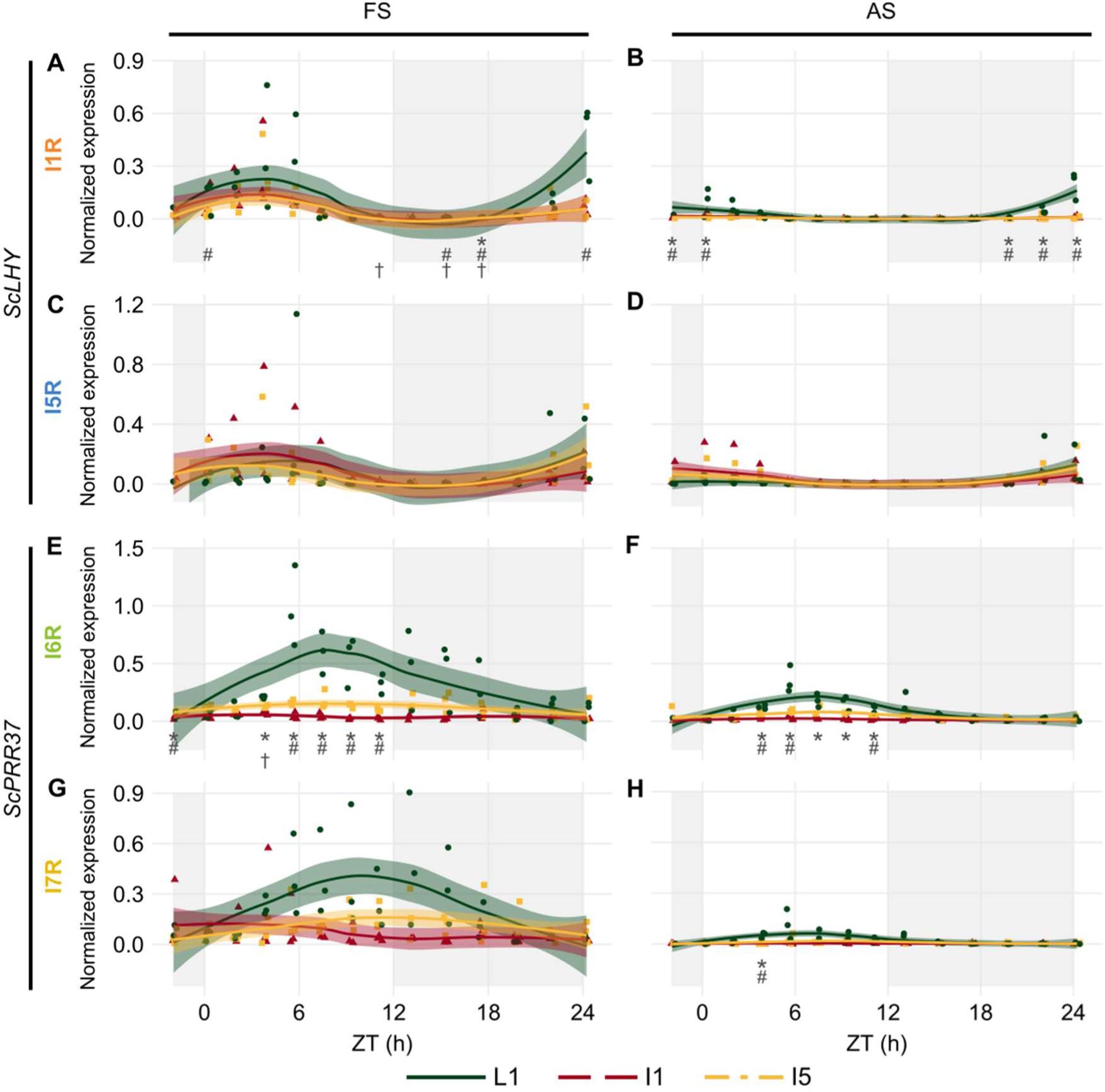
Diel expression profile of fully spliced and alternative spliced transcript isoforms in leaves and internodes. Biological replicates (circles and triangles) and their LOESS curve (continuous lines ± SE) of fully spliced (left column) and alternative spliced (right column) transcript forms of sugarcane circadian clock genes during the summer harvest in leaves (green), internode 1 and 2 (red) and internode 5 (yellow). (A, B) *ScLHY* gene expression shows levels of I1R and **(C, D)** I5R; (E, F) *ScPRR37* gene expression shows levels of I6R and **(G, H)** I7R. The light-gray boxes represent the night period. Statistical significance was analyzed by One-way ANOVA with post-hoc Tukey HSD test, * p < 0.05 when comparing L1 against I1, # p < 0.05 when comparing L1 against I5, and † p < 0.05 when comparing I1 against I5.

**Table S1.**
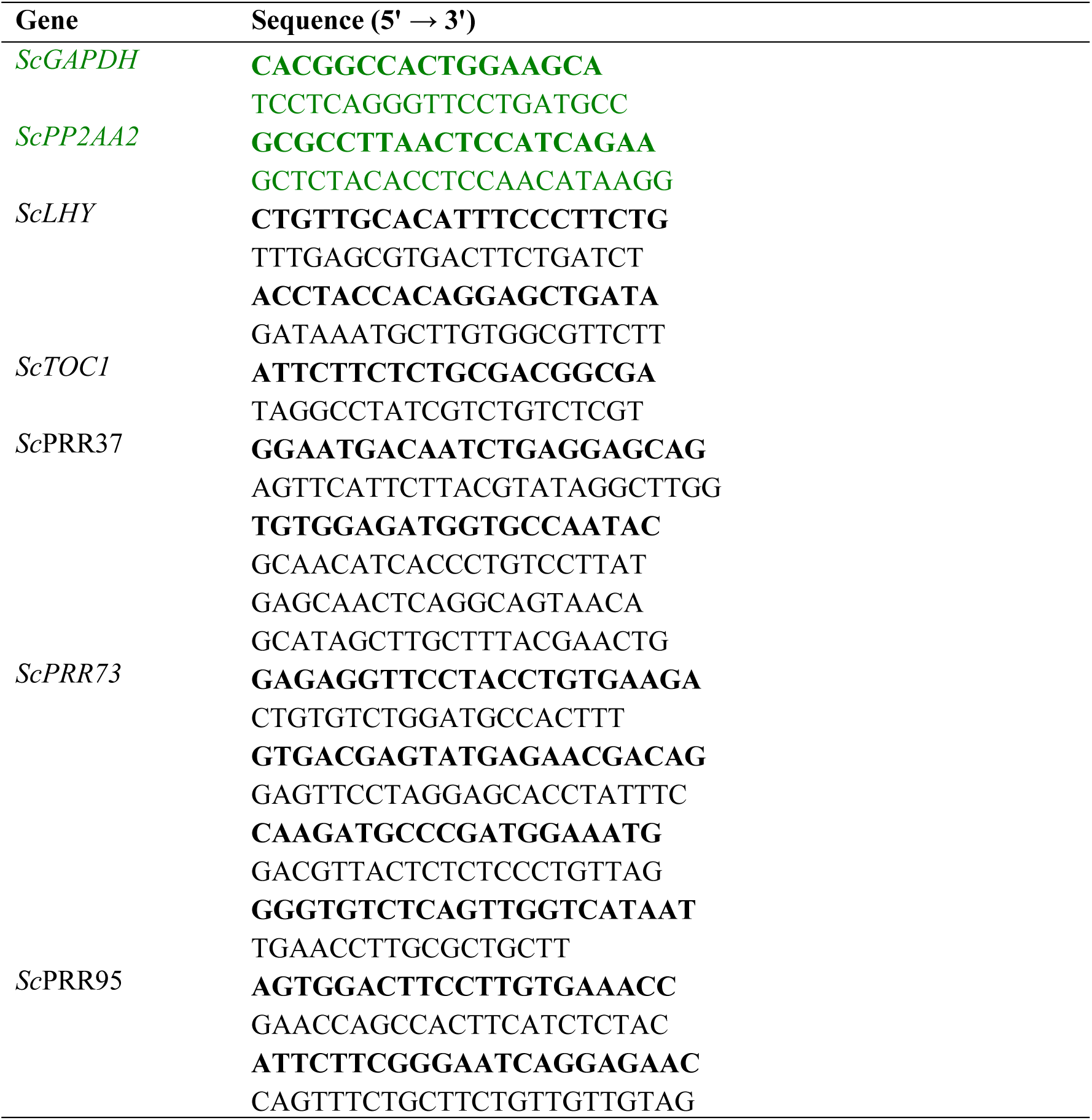
List of sugarcane gene-specific primers and their sequences. Primers in bold indicate forward (FW) sequences, as well as the primers bearing the 6-FAM tag used in the HR RT-PCR assays. The other sequences indicate reverse (RV) sequences. Normalizer primers are highlighted in green.

**TableS2.**
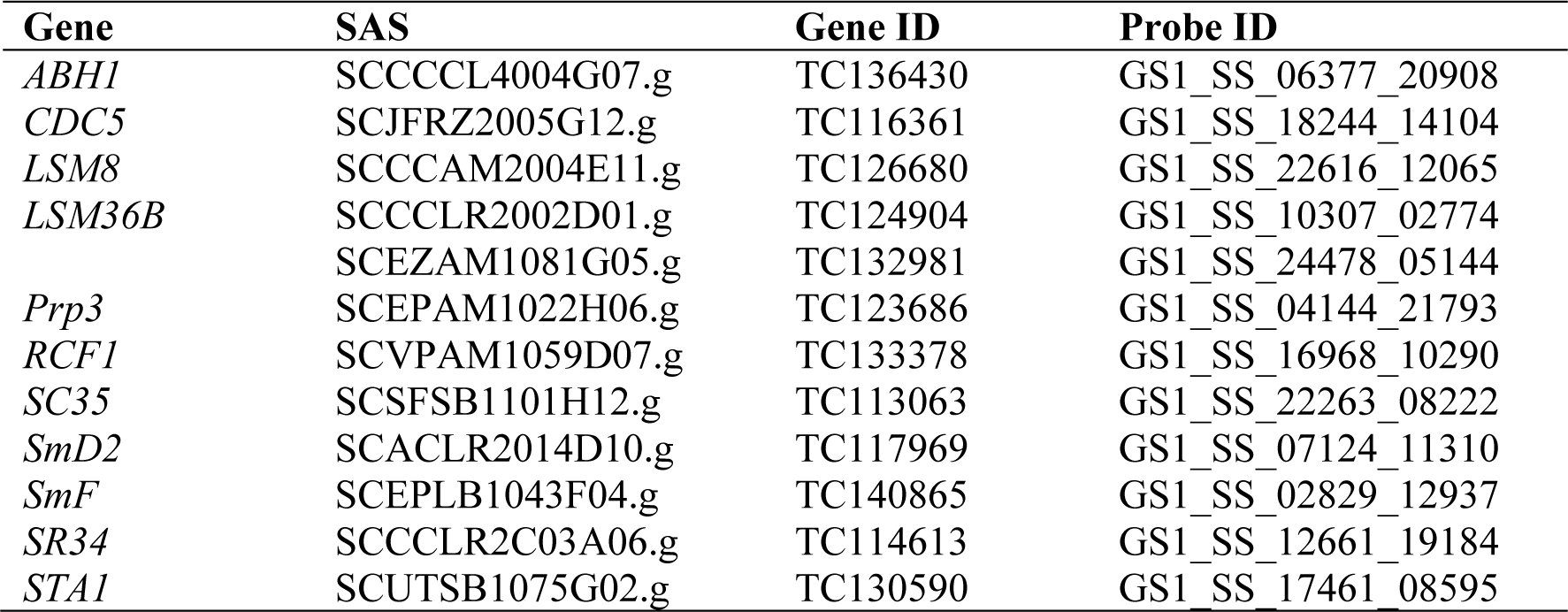
SAS number, GeneID, and ProbeID of genes associated with the Spliceosome in the Sugarcane array (Dantas et al., 2019).

**Table S3.**
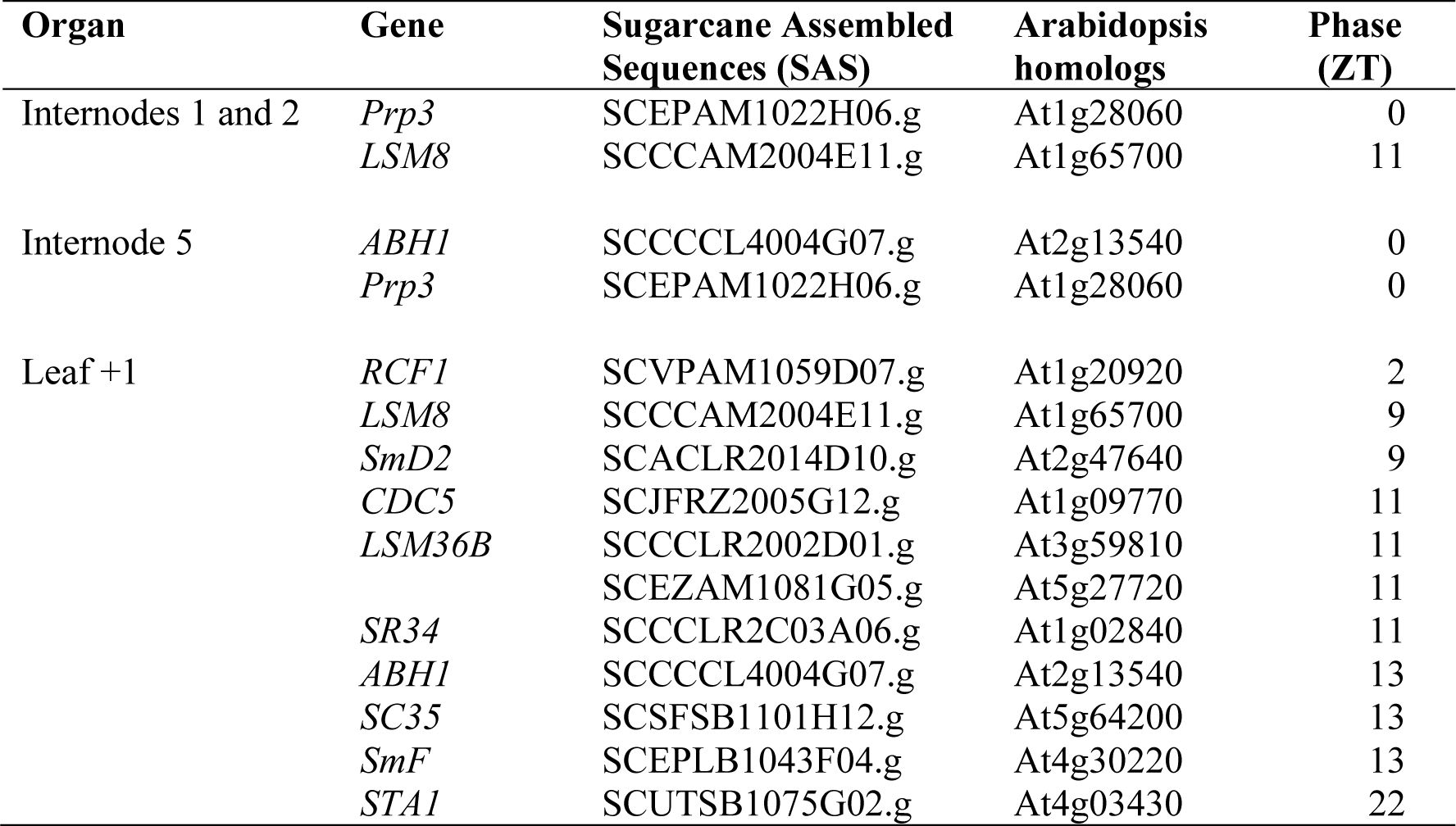
Spliceosome-related transcripts showing rhythmicity in field conditions according to Dantas et al. (2019). Sugarcane Assembled Sequences (SAS) were taken from the SUCEST-FUN database (http://sucest-fun.org).

## References

Alabadí, D., Oyama, T., Yanovsky, M. J., Harmon, F. G., Más, P., and Kay, S. A. (2001). Reciprocal Regulation Between TOC1 and LHY / CCA1 Within the Arabidopsis Circadian Clock. Science 293, 880–883. doi:10.1126/science.1061320.

Amorim, H. V., Lopes, M. L., De Castro Oliveira, J. V., Buckeridge, M. S., and Goldman, G. H. (2011). Scientific challenges of bioethanol production in Brazil. Applied Microbiology and Biotechnology 91, 1267–1275. doi:10.1007/s00253-011-3437-6.

Annunziata, M. G., Apelt, F., Carillo, P., Krause, U., Feil, R., Koehl, K., et al. (2018). Response of Arabidopsis primary metabolism and circadian clock to low night temperature in a natural light environment. Journal of Experimental Botany 69, 4881–4895. doi:10.1093/jxb/ery276.

Annunziata, M. G., Apelt, F., Carillo, P., Krause, U., Feil, R., Mengin, V., et al. (2017). Getting back to nature: a reality check for experiments in controlled environments. Journal of Experimental Botany 68, 4463–4477. doi:10.1093/jxb/erx220.

Beyer, A. L., and Osheim, Y. N. (1988). Splice site selection, rate of splicing, and alternative splicing on nascent transcripts. Genes & development 2, 754–765. doi:10.1101/gad.2.6.754.

Bieniawska, Z., Espinoza, C., Schlereth, A., Sulpice, R., Hincha, D. K., and Hannah, M. A. (2008). Disruption of the Arabidopsis Circadian Clock Is Responsible for Extensive Variation in the. 147, 263–279. doi:10.1104/pp.108.118059.

Calixto, C. P. G., Guo, W., James, A. B., Tzioutziou, N. A., Entizne, J. C., Panter, P. E., et al. (2018). Rapid and Dynamic Alternative Splicing Impacts the Arabidopsis Cold Response Transcriptome. Plant Cell 30, 1424–1444. doi:10.1105/tpc.18.00177.

Calixto, C. P. G., Simpson, C. G., Waugh, R., and Brown, J. W. S. (2016). Alternative Splicing of Barley Clock Genes in Response to Low Temperature. PLOS ONE 11, e0168028. doi:10.1371/journal.pone.0168028.

Calixto, C. P. G., Tzioutziou, N. A., James, A. B., Hornyik, C., Guo, W., Zhang, R., et al. (2019). Cold-Dependent Expression and Alternative Splicing of Arabidopsis Long Non-coding RNAs. Front Plant Sci 10, 235. doi:10.3389/fpls.2019.00235.

Calixto, C. P. G., Waugh, R., and Brown, J. W. S. (2015). Evolutionary Relationships Among Barley and Arabidopsis Core Circadian Clock and Clock-Associated Genes. Journal of Molecular Evolution 80, 108–119. doi:10.1007/s00239-015-9665-0.

Chamala, S., Feng, G., Chavarro, C., and Barbazuk, W. B. (2015). Genome-wide identification of evolutionarily conserved alternative splicing events in flowering plants. Front Bioeng Biotechnol 3, 33. doi:10.3389/fbioe.2015.00033.

Chan, A., Charron, C., van der Vossen, E., D’Hont, A., Town, C., Hervouet, C., et al. (2018). A mosaic monoploid reference sequence for the highly complex genome of sugarcane. Nature Communications 9. doi:10.1038/s41467-018-05051-5.

Chaudhary, S., Jabre, I., Reddy, A. S. N., Staiger, D., and Syed, N. H. (2019). Perspective on Alternative Splicing and Proteome Complexity in Plants. Trends in Plant Science 0. doi:10.1016/j.tplants.2019.02.006.

Cuadrado, A., Acevedo, R., Moreno Díaz De La Espina, S., Jouve, N., and De La Torre, C. (2004). Genome remodelling in three modern S. officinarumxS. spontaneum sugarcane cultivars. Journal of Experimental Botany 55, 847–854. doi:10.1093/jxb/erh093.

Dantas, L. L. B., de Lima, N. O., Nishiyama, M. Y., Carneiro, M. S., Souza, G. M., and Hotta, C. T. (2019). Rhythms of Transcription in Field-Grown Sugarcane Are Highly Organ Specific. BioRXvs. doi:https://doi.org/10.1101/607002.

de Setta, N., Monteiro-Vitorello, C., Metcalfe, C., Cruz, G. M., Del Bem, L., Vicentini, R., et al. (2014). Building the sugarcane genome for biotechnology and identifying evolutionary trends. BMC Genomics 15, 540. doi:10.1186/1471-2164-15-540.

D’Hont, A. (2005). Unraveling the genome structure of polyploids using FISH and GISH; examples of sugarcane and banana. Cytogenetic and Genome Research 109, 27–33. doi:10.1159/000082378.

D’Hont, A., Grivet, L., Feldmann, P., Rao, S., Berding, N., and Glaszmann, J. C. (1996). Characterization of the double genome structure of sugarcane cultivars (Saccharum ssp.) by molecular cytogenetics. Mol. Gen. Genet. 250, 405–413.

Ding, F., Cui, P., Wang, Z., Zhang, S., Ali, S., and Xiong, L. (2014). Genome-wide analysis of alternative splicing of pre-mRNA under salt stress in Arabidopsis. BMC genomics 15, 431. doi:10.1186/1471-2164-15-431.

Dodd, A. N. (2009). Plant Circadian Clocks Increase Photosynthesis, Growth, Survival,. Science 630, 630–3. doi:10.1126/science.1115581.

Dodd, A. N., Love, J., and Webb, A. A. R. (2005). The plant clock shows its metal: Circadian regulation of cytosolic free Ca 2+. Trends in Plant Science 10, 15–21. doi:10.1016/j.tplants.2004.12.001.

Endo, M., Shimizu, H., Nohales, M. a., Araki, T., and Kay, S. a. (2014). Tissue-specific clocks in Arabidopsis show asymmetric coupling. Nature 515, 419–422. doi:10.1038/nature13919.

Filichkin, S. A., Cumbie, J. S., Dharmawardhana, P., Jaiswal, P., Chang, J. H., Palusa, S. G., et al. (2015a). Environmental Stresses Modulate Abundance and Timing of Alternatively Spliced Circadian Transcripts in Arabidopsis. Molecular plant 8, 207–27. doi:10.1016/j.molp.2014.10.011.

Filichkin, S. A., Cumbie, J. S., Dharmawardhana, P., Jaiswal, P., Chang, J. H., Palusa, S. G., et al. (2015b). Environmental stresses modulate abundance and timing of alternatively spliced circadian transcripts in Arabidopsis. Molecular Plant 8, 207–227. doi:10.1016/j.molp.2014.10.011.

Filichkin, S. a, and Mockler, T. C. (2012). Unproductive alternative splicing and nonsense mRNAs: A widespread phenomenon among plant circadian clock genes. Biology Direct 7, 20. doi:10.1186/1745-6150-7-20.

Filichkin, S. A., Priest, H. D., Givan, S. A., Shen, R., Bryant, D. W., Fox, S. E., et al. (2010). Genome-wide mapping of alternative splicing in Arabidopsis thaliana. 45–58. doi:10.1101/gr.093302.109.2008.

Filichkin, S., Priest, H. D., Megraw, M., and Mockler, T. C. (2015c). Alternative splicing in plants: Directing traffic at the crossroads of adaptation and environmental stress. Current Opinion in Plant Biology 24, 125–135. doi:10.1016/j.pbi.2015.02.008.

Fouquet, R., Martin, F., Fajardo, D. S., Gault, C. M., Gómez, E., Tseung, C.-W., et al. (2011). Maize rough endosperm3 encodes an RNA splicing factor required for endosperm cell differentiation and has a nonautonomous effect on embryo development. Plant Cell 23, 4280–97. doi:10.1105/tpc.111.092163.

Frank, A., Matiolli, C. C., Viana, A. J. C., Hearn, T. J., Kusakina, J., Belbin, F. E., et al. (2018). Circadian entrainment in Arabidopsis by the sugar-responsive transcription factor bZIP63. Current Biology 28, 2597–2606.e6. doi:10.1016/j.cub.2018.05.092.

Fung-Uceda, J., Lee, K., Seo, P. J., Polyn, S., De Veylder, L., and Mas, P. (2018). The Circadian Clock Sets the Time of DNA Replication Licensing to Regulate Growth in Arabidopsis. Dev. Cell 45, 101–113.e4. doi:10.1016/j.devcel.2018.02.022.

Garcia, A. a F., Mollinari, M., Marconi, T. G., Serang, O. R., Silva, R. R., Vieira, M. L. C., et al. (2013). SNP genotyping allows an in-depth characterisation of the genome of sugarcane and other complex autopolyploids. Scientific reports 3, 3399. doi:10.1038/srep03399.

Garsmeur, O., Droc, G., Antonise, R., Grimwood, J., Potier, B., Aitken, K., et al. (2018). A mosaic monoploid reference sequence for the highly complex genome of sugarcane. Nat Commun 9, 2638. doi:10.1038/s41467-018-05051-5.

Godoy Herz, M. A., Kubaczka, M. G., Brzyżek, G., Servi, L., Krzyszton, M., Simpson, C., et al. (2019). Light Regulates Plant Alternative Splicing through the Control of Transcriptional Elongation. Molecular Cell, 1066–1074. doi:10.1016/j.molcel.2018.12.005.

Göhring, J., Jacak, J., and Barta, A. (2014). Imaging of endogenous messenger RNA splice variants in living cells reveals nuclear retention of transcripts inaccessible to nonsense-mediated decay in Arabidopsis. Plant Cell 26, 754–764. doi:10.1105/tpc.113.118075.

Graf, A., Schlereth, A., Stitt, M., and Smith, A. M. (2010). Circadian control of carbohydrate availability for growth in Arabidopsis plants at night. Proc. Natl. Acad. Sci. U.S.A. 107, 9458–9463. doi:10.1073/pnas.0914299107.

Graf, A., and Smith, A. M. (2011). Starch and the clock: the dark side of plant productivity. Trends Plant Sci. 16, 169–175. doi:10.1016/j.tplants.2010.12.003.

Graveley, B. R. (2005). Mutually exclusive splicing of the insect Dscam Pre-mRNA directed by competing intronic RNA secondary structures. Cell 123, 65–73. doi:10.1016/j.cell.2005.07.028.

Grundy, J., Stoker, C., and Carré, I. A. (2015). Circadian regulation of abiotic stress tolerance in plants. Frontiers in Plant Science 6, 648. doi:10.3389/fpls.2015.00648.

Harmer, S. L. (2009). The circadian system in higher plants. Annual review of plant biology 60, 357–77. doi:10.1146/annurev.arplant.043008.092054.

Haydon, M. J., Mielczarek, O., Robertson, F. C., Hubbard, K. E., and Webb, A. A. R. (2013). Photosynthetic entrainment of the Arabidopsis thaliana circadian clock. Nature 502, 689–92. doi:10.1038/nature12603.

Higashi, T., Tanigaki, Y., Takayama, K., Nagano, A. J., Honjo, M. N., and Fukuda, H. (2016). Detection of diurnal variation of tomato transcriptome through the molecular timetable method in a sunlight-type plant factory. Frontiers in Plant Science 7, 1–9. doi:10.3389/fpls.2016.00087.

Hotta, C. T., Nishiyama, M. Y., and Souza, G. M. (2013). Circadian Rhythms of Sense and Antisense Transcription in Sugarcane, a Highly Polyploid Crop. PLoS ONE 8. doi:10.1371/journal.pone.0071847.

Hsu, P. Y., and Harmer, S. L. (2014). Wheels within wheels: The plant circadian system. Trends in Plant Science 19, 240–249. doi:10.1016/j.tplants.2013.11.007.

Iskandar, H. M., Simpson, R. S., Casu, R. E., Bonnett, G. D., Maclean, D. J., and Manners, J. M. (2004). Comparison of reference genes for quantitative real-time polymerase chain reaction analysis of gene expression in sugarcane. Plant Mol Biol Rep 22, 325–337. doi:10.1007/BF02772676.

Izawa, T., Mihara, M., Suzuki, Y., Gupta, M., Itoh, H., Nagano, A. J., et al. (2011a). Os-GIGANTEA Confers Robust Diurnal Rhythms on the Global Transcriptome of Rice in the Field. Plant Cell 23, 1741–1755. doi:10.1105/tpc.111.083238.

Izawa, T., Mihara, M., Suzuki, Y., Gupta, M., Itoh, H., Nagano, A. J., et al. (2011b). Os-GIGANTEA confers robust diurnal rhythms on the global transcriptome of rice in the field. Plant Cell 23, 1741–1755. doi:10.1105/tpc.111.083238.

Jabre, I., Reddy, A. S. N., Kalyna, M., Chaudhary, S., Khokhar, W., Byrne, L. J., et al. (2019). Does co-transcriptional regulation of alternative splicing mediate plant stress responses? Nucleic Acids Res 47, 2716–2726. doi:10.1093/nar/gkz121.

Jacob, A. G., and Smith, C. W. J. (2017). Intron retention as a component of regulated gene expression programs. Hum. Genet. 136, 1043–1057. doi:10.1007/s00439-017-1791-x.

James, A. B., Syed, N. H., Bordage, S., Marshall, J., Nimmo, G. A., Jenkins, G. I., et al. (2012a). Alternative splicing mediates responses of the Arabidopsis circadian clock to temperature changes. Plant Cell 24, 961–981. doi:10.1105/tpc.111.093948.

James, A. B., Syed, N. H., Bordage, S., Marshall, J., Nimmo, G. A., Jenkins, G. I., et al. (2012b). Alternative Splicing Mediates Responses of the Arabidopsis Circadian Clock to Temperature Changes. The Plant Cell 24, 961–981. doi:10.1105/tpc.111.093948.

James, A. B., Syed, N. H., Brown, J. W. S., and Nimmo, H. G. (2012c). Thermoplasticity in the plant circadian clock: how plants tell the time-perature. Plant signaling & behavior 7, 1219–23. doi:10.4161/psb.21491.

Jones, M. A., Williams, B. A., McNicol, J., Simpson, C. G., Brown, J. W. S., and Harmer, S. L. (2012). Mutation of *Arabidopsis SPLICEOSOMAL TIMEKEEPER LOCUS1* Causes Circadian Clock Defects. The Plant Cell 24, 4066–4082. doi:10.1105/tpc.112.104828.

Kalyna, M., Simpson, C. G., Syed, N. H., Lewandowska, D., Marquez, Y., Kusenda, B., et al. (2012). Alternative splicing and nonsense-mediated decay modulate expression of important regulatory genes in Arabidopsis. Nucleic Acids Research 40, 2454–2469. doi:10.1093/nar/gkr932.

Khan, S., Rowe, S. C., and Harmon, F. G. (2010). Coordination of the maize transcriptome by a conserved circadian clock. BMC plant biology 10, 126. doi:10.1186/1471-2229-10-126.

Ko, D. K., Rohozinski, D., Song, Q., Taylor, S. H., Juenger, T. E., Harmon, F. G., et al. (2016). Temporal Shift of Circadian-Mediated Gene Expression and Carbon Fixation Contributes to Biomass Heterosis in Maize Hybrids. PLoS Genet. 12, e1006197. doi:10.1371/journal.pgen.1006197.

Kornblihtt, A. R., Schor, I. E., Alló, M., Dujardin, G., Petrillo, E., and Muñoz, M. J. (2013). Alternative splicing: a pivotal step between eukaryotic transcription and translation. Nature reviews. Molecular cell biology 14, 153–65. doi:10.1038/nrm3525.

Kriechbaumer, V., Wang, P., Hawes, C., and Abell, B. M. (2012). Alternative splicing of the auxin biosynthesis gene YUCCA4 determines its subcellular compartmentation. Plant Journal 70, 292–302. doi:10.1111/j.1365-313X.2011.04866.x.

Kwon, Y.-J., Park, M.-J., Kim, S.-G., Baldwin, I. T., and Park, C.-M. (2014). Alternative splicing and nonsense-mediated decay of circadian clock genes under environmental stress conditions in Arabidopsis. BMC Plant Biol. 14, 136. doi:10.1186/1471-2229-14-136.

Lai, A. G., Doherty, C. J., Mueller-Roeber, B., Kay, S. A., Schippers, J. H. M., and Dijkwel, P. P. (2012). CIRCADIAN CLOCK-ASSOCIATED 1 regulates ROS homeostasis and oxidative stress responses. Proceedings of the National Academy of Sciences 109, 17129–17134. doi:10.1073/pnas.1209148109.

Liu, M., Yuan, L., Liu, N. Y., Shi, D. Q., Liu, J., and Yang, W. C. (2009). GAMETOPHYTIC FACTOR 1, involved in pre-mRNA splicing, is essential for megagametogenesis and embryogenesis in Arabidopsis. Journal of Integrative Plant Biology 51, 261–271. doi:10.1111/j.1744-7909.2008.00783.x.

Locke, J. C. W., Millar, A. J., and Turner, M. S. (2005). Modelling genetic networks with noisy and varied experimental data: The circadian clock in Arabidopsis thaliana. Journal of Theoretical Biology 234, 383–393. doi:10.1016/j.jtbi.2004.11.038.

Lu, Y., Gehan, J. P., and Sharkey, T. D. (2005). Daylength and circadian effects on starch degradation and maltose metabolism. Plant physiology 138, 2280–2291. doi:10.1104/pp.105.061903.

Marquez, Y., Brown, J. W. S., Simpson, C., Barta, A., and Kalyna, M. (2012). Transcriptome survey reveals increased complexity of the alternative splicing landscape in Arabidopsis. Genome Research 22, 1184–1195. doi:10.1101/gr.134106.111.

Marshall, C. M., Tartaglio, V., Duarte, M., and Harmon, F. G. (2016). The Arabidopsis sickle Mutant Exhibits Altered Circadian Clock Responses to Cool Temperatures and Temperature-Dependent Alternative Splicing. Plant Cell 28, 2560–2575. doi:10.1105/tpc.16.00223.

Mastrangelo, A. M., Marone, D., Laidò, G., De Leonardis, A. M., and De Vita, P. (2012). Alternative splicing: Enhancing ability to cope with stress via transcriptome plasticity. Plant Science 185–186, 40–49. doi:10.1016/j.plantsci.2011.09.006.

McClung, C. (2019). The Plant Circadian Oscillator. Biology 8, 14. doi:10.3390/biology8010014.

McCormick, R. F., Truong, S. K., Sreedasyam, A., Jenkins, J., Shu, S., Sims, D., et al. (2018). The Sorghum bicolor reference genome: improved assembly, gene annotations, a transcriptome atlas, and signatures of genome organization. Plant J. 93, 338–354. doi:10.1111/tpj.13781.

Millar, A. J. (2016). The Intracellular Dynamics of Circadian Clocks Reach for the Light of Ecology and Evolution. Annual Review of Plant Biology 67, annurev-arplant-043014-115619. doi:10.1146/annurev-arplant-043014-115619.

Miller, M., Zhang, C., and Chen, Z. J. (2012). Ploidy and Hybridity Effects on Growth Vigor and Gene Expression in Arabidopsis thaliana Hybrids and Their Parents. G3 (Bethesda) 2, 505–513. doi:10.1534/g3.112.002162.

Min, X. J., Powell, B., Braessler, J., Meinken, J., Yu, F., and Sablok, G. (2015). Genome-wide cataloging and analysis of alternatively spliced genes in cereal crops. BMC genomics 16, 721. doi:10.1186/s12864-015-1914-5.

Moll, C., Von Lyncker, L., Zimmermann, S., Kägi, C., Baumann, N., Twell, D., et al. (2008). CLO/GFA1 and ATO are novel regulators of gametic cell fate in plants. Plant Journal 56, 913–921. doi:10.1111/j.1365-313X.2008.03650.x.

Moore, P. (1995). Temporal and Spatial Regulation of Sucrose Accumulation in the Sugarcane Stem. Australian Journal of Plant Physiology 22, 661. doi:10.1071/PP9950661.

Murakami, M., Tago, Y., Yamashino, T., and Mizuno, T. (2007). Comparative overviews of clock-associated genes of Arabidopsis thaliana and Oryza sativa. Plant and Cell Physiology 48, 110–121. doi:10.1093/pcp/pcl043.

Nagano, A. J., Sato, Y., Mihara, M., Antonio, B. A., Motoyama, R., Itoh, H., et al. (2012). Deciphering and prediction of transcriptome dynamics under fluctuating field conditions. Cell 151, 1358–1369. doi:10.1016/j.cell.2012.10.048.

Nagashima, Y., Mishiba, K.-I., Suzuki, E., Shimada, Y., Iwata, Y., and Koizumi, N. (2011). Arabidopsis IRE1 catalyses unconventional splicing of bZIP60 mRNA to produce the active transcription factor. Scientific reports 1, 29. doi:10.1038/srep00029.

Nakamichi, N., Kiba, T., Henriques, R., Mizuno, T., Chua, N.-H., and Sakakibara, H. (2010). PSEUDO-RESPONSE REGULATORS 9, 7, and 5 are transcriptional repressors in the Arabidopsis circadian clock. The Plant cell 22, 594–605. doi:10.1105/tpc.109.072892.

Ng, D. W.-K., Miller, M., Yu, H. H., Huang, T.-Y., Kim, E.-D., Lu, J., et al. (2014). A Role for CHH Methylation in the Parent-of-Origin Effect on Altered Circadian Rhythms and Biomass Heterosis in Arabidopsis Intraspecific Hybrids. Plant Cell 26, 2430–2440. doi:10.1105/tpc.113.115980.

Ni, Z., Kim, E.-D., Ha, M., Lackey, E., Liu, J., Zhang, Y., et al. (2009). Altered circadian rhythms regulate growth vigour in hybrids and allopolyploids. Nature 457, 327–331. doi:10.1038/nature07523.

Oesterreich, F. C., Herzel, L., Strauber, K., Hujer, K., Howard, J., and Neugebauer, K. M. (2016). Splicing of Nascent RNA Coincides with Intron Exit from RNA Polymerase II. Cell 165, 372–381. doi:10.1016/j.cell.2016.02.045.

Palusa, S. G., Ali, G. S., and Reddy, A. S. N. (2007). Alternative splicing of pre-mRNAs of Arabidopsis serine/arginine-rich proteins: Regulation by hormones and stresses. Plant Journal 49, 1091–1107. doi:10.1111/j.1365-313X.2006.03020.x.

Para, A., Farré, E. M., Imaizumi, T., Pruneda-Paz, J. L., Harmon, F. G., and Kay, S. A. (2007). PRR3 Is a vascular regulator of TOC1 stability in the Arabidopsis circadian clock. The Plant cell 19, 3462–73. doi:10.1105/tpc.107.054775.

Park, M.-J., Seo, P. J., and Park, C.-M. (2012). CCA1 alternative splicing as a way of linking the circadian clock to temperature response in Arabidopsis. Plant Signal Behav 7, 1194–1196. doi:10.4161/psb.21300.

Pokhilko, A., Fernández, A. P., Edwards, K. D., Southern, M. M., Halliday, K. J., and Millar, A. J. (2012). The clock gene circuit in Arabidopsis includes a repressilator with additional feedback loops. Molecular systems biology 8, 574. doi:10.1038/msb.2012.6.

Reddy, A. S. N., Marquez, Y., Kalyna, M., and Barta, A. (2013). Complexity of the alternative splicing landscape in plants. The Plant cell 25, 3657–83. doi:10.1105/tpc.113.117523.

Remy, E., Cabrito, T. R., Baster, P., Batista, R. A., Teixeira, M. C., Friml, J., et al. (2013). A major facilitator superfamily transporter plays a dual role in polar auxin transport and drought stress tolerance in Arabidopsis. The Plant cell 25, 901–26. doi:10.1105/tpc.113.110353.

Riaño-Pachón, D. M., and Mattiello, L. (2017). Draft genome sequencing of the sugarcane hybrid SP80-3280. F1000Res 6, 861. doi:10.12688/f1000research.11859.2.

Richards, C. L., Rosas, U., Banta, J., Bhambhra, N., and Purugganan, M. D. (2012). Genome-wide patterns of Arabidopsis gene expression in nature. PLoS Genetics 8. doi:10.1371/journal.pgen.1002662.

Rosloski, S. M., Singh, A., Jali, S. S., Balasubramanian, S., Weigel, D., and Grbic, V. (2013). Functional analysis of splice variant expression of MADS AFFECTING FLOWERING 2 of Arabidopsis thaliana. Plant Molecular Biology 81, 57–69. doi:10.1007/s11103-012-9982-2.

Saldi, T., Cortazar, M. A., Sheridan, R. M., and Bentley, D. L. (2016). Coupling of RNA Polymerase II Transcription Elongation with Pre-mRNA Splicing. Journal of Molecular Biology 428, 2623–2635. doi:10.1016/j.jmb.2016.04.017.

Sato, Y., Antonio, B., Namiki, N., Motoyama, R., Sugimoto, K., Takehisa, H., et al. (2011). Field transcriptome revealed critical developmental and physiological transitions involved in the expression of growth potential in japonica rice. BMC plant biology 11, 10. doi:10.1186/1471-2229-11-10.

Seo, P. J., Hong, S. Y., Kim, S. G., and Park, C. M. (2011). Competitive inhibition of transcription factors by small interfering peptides. Trends in Plant Science 16, 541–549. doi:10.1016/j.tplants.2011.06.001.

Seo, P. J., Park, M.-J., Lim, M.-H., Kim, S.-G., Lee, M., Baldwin, I. T., et al. (2012). A self-regulatory circuit of CIRCADIAN CLOCK-ASSOCIATED1 underlies the circadian clock regulation of temperature responses in Arabidopsis. Plant Cell 24, 2427–2442. doi:10.1105/tpc.112.098723.

Severing, E. I., van Dijk, A. D. J., Morabito, G., Busscher-Lange, J., Immink, R. G. H., and van Ham, R. C. H. J. (2012). Predicting the impact of alternative splicing on plant MADS domain protein function. PLoS ONE 7. doi:10.1371/journal.pone.0030524.

Shalit-Kaneh, A., Kumimoto, R. W., Filkov, V., and Harmer, S. L. (2018). Multiple feedback loops of the Arabidopsis circadian clock provide rhythmic robustness across environmental conditions. Proc. Natl. Acad. Sci. U.S.A. 115, 7147–7152. doi:10.1073/pnas.1805524115.

Shang, X., Cao, Y., and Ma, L. (2017a). Alternative splicing in plant genes: A means of regulating the environmental fitness of plants. International Journal of Molecular Sciences 18. doi:10.3390/ijms18020432.

Shang, X., Cao, Y., and Ma, L. (2017b). Alternative Splicing in Plant Genes: A Means of Regulating the Environmental Fitness of Plants. Int J Mol Sci 18. doi:10.3390/ijms18020432.

Shen, Y., Zhou, Z., Wang, Z., Li, W., Fang, C., Wu, M., et al. (2014). Global Dissection of Alternative Splicing in Paleopolyploid Soybean. The Plant Cell 26, 996–1008. doi:10.1105/tpc.114.122739.

Simpson, C. G., Fuller, J., Maronova, M., Kalyna, M., Davidson, D., McNicol, J., et al. (2007). Monitoring changes in alternative precursor messenger RNA splicing in multiple gene transcripts. The Plant Journal 53, 1035–1048. doi:10.1111/j.1365-313X.2007.03392.x.

Simpson, C. G., Fuller, J., Rapazote-Flores, P., Mayer, C. D., Calixto, C. P. G., Milne, L., et al. (2019). High-resolution RT-PCR analysis of alternative barley transcripts. Methods in Molecular Biology 1900, 269–281. doi:10.1007/978-1-4939-8944-7_17.

Souza, G. M., Sluys, M. A. V., Lembke, C. G., Lee, H., Margarido, G. R. A., Hotta, C. T., et al. (unpublished). Assembly of the 373K gene space of the polyploid sugarcane genome reveals reservoirs of functional diversity in the world’s leading biomass crop.

Staiger, D., and Brown, J. W. S. (2013a). Alternative splicing at the intersection of biological timing, development, and stress responses. Plant Cell 25, 3640–3656. doi:10.1105/tpc.113.113803.

Staiger, D., and Brown, J. W. S. (2013b). Alternative Splicing at the Intersection of Biological Timing, Development, and Stress Responses. The Plant Cell 25, 3640–3656. doi:10.1105/tpc.113.113803.

Sugliani, M., Brambilla, V., Clerkx, E. J. M., Koornneef, M., and Soppe, W. J. J. (2010). The conserved splicing factor SUA controls alternative splicing of the developmental regulator ABI3 in Arabidopsis. The Plant cell 22, 1936–46. doi:10.1105/tpc.110.074674.

Syed, N. H., Kalyna, M., Marquez, Y., Barta, A., and Brown, J. W. S. (2012). Alternative splicing in plants – coming of age. Trends in Plant Science 17, 616–623. doi:10.1016/j.tplants.2012.06.001.

Szakonyi, D., and Duque, P. (2018). Alternative Splicing as a Regulator of Early Plant Development. Frontiers in Plant Science 9, 1–9. doi:10.3389/fpls.2018.01174.

Thatcher, S. R., Zhou, W., Leonard, A., Wang, B.-B., Beatty, M., Zastrow-Hayes, G., et al. (2014). Genome-Wide Analysis of Alternative Splicing in Zea mays: Landscape and Genetic Regulation. The Plant Cell Online 26, 3472–3487. doi:10.1105/tpc.114.130773.

Vaneechoutte, D., Estrada, A. R., Lin, Y. C., Loraine, A. E., and Vandepoele, K. (2017). Genome-wide characterization of differential transcript usage in Arabidopsis thaliana. Plant Journal 92, 1218–1231. doi:10.1111/tpj.13746.

Vicentini, R., Del Bem, L. E. V., Van Sluys, M. A., Nogueira, F. T. S., and Vincentz, M. (2012). Gene Content Analysis of Sugarcane Public ESTs Reveals Thousands of Missing Coding-Genes and an Unexpected Pool of Grasses Conserved ncRNAs. Tropical Plant Biol. 5, 199–205. doi:10.1007/s12042-012-9103-z.

Wang, X., Yang, M., Ren, D., Terzaghi, W., Deng, X. W., and He, G. (2018). Cis-regulated alternative splicing divergence and its potential contribution to environmental responses in Arabidopsis. Plant Journal, 555–570. doi:10.1111/tpj.14142.

Zeilinger, M. N., Farré, E. M., Taylor, S. R., Kay, S. a, and Doyle, F. J. (2006). A novel computational model of the circadian clock in Arabidopsis that incorporates PRR7 and PRR9. Mol. Syst. Biol. 2, 58. doi:10.1038/msb4100101.

Zhang, J., Zhang, X., Tang, H., Zhang, Q., Hua, X., Ma, X., et al. (2018). Allele-defined genome of the autopolyploid sugarcane Saccharum spontaneum L. Nature Genetics 50, 1565. doi:10.1038/s41588-018-0237-2.

Zhang, Q., Zhang, X., Wang, S., Tan, C., Zhou, G., and Li, C. (2016). Involvement of Alternative Splicing in Barley Seed Germination. PloS one 11, e0152824. doi:10.1371/journal.pone.0152824.

Zhang, X., Fang, X., Li, R., Li, S., Zhang, G., Tao, Y., et al. (2010). Deep RNA sequencing at single base-pair resolution reveals high complexity of the rice transcriptome. Genome Research 20, 646–654. doi:10.1101/gr.100677.109.

Zhang, Z., Zhang, S., Zhang, Y., Wang, X., Li, D., Li, Q., et al. (2011). Arabidopsis floral initiator SKB1 confers high salt tolerance by regulating transcription and pre-mRNA splicing through altering histone H4R3 and small nuclear ribonucleoprotein LSM4 methylation. The Plant cell 23, 396–411. doi:10.1105/tpc.110.081356.

